# Persistent delay in maturation of the developing gut microbiota in infants with cystic fibrosis

**DOI:** 10.1101/2023.05.02.539134

**Authors:** Paige Salerno, Adrian Verster, Rebecca Valls, Kaitlyn Barrack, Courtney Price, Juliette Madan, George A. O’Toole, Benjamin D. Ross

## Abstract

The healthy human infant gut microbiome undergoes stereotypical changes in taxonomic composition between birth and maturation to an adult-like stable state. During this time, extensive communication between microbiota and the host immune system contributes to health status later in life. Although there are many reported associations between microbiota compositional alterations and disease in adults, less is known about how microbiome development is altered in pediatric diseases. One pediatric disease linked to altered gut microbiota composition is cystic fibrosis (CF), a multi-organ genetic disease involving impaired chloride secretion across epithelia and heightened inflammation both in the gut and at other body sites. Here, we use shotgun metagenomics to profile the strain-level composition and developmental dynamics of the infant fecal microbiota from several CF and non-CF longitudinal cohorts spanning from birth to greater than 36 months of life. We identify a set of keystone species whose prevalence and abundance reproducibly define microbiota development in early life in non-CF infants, but are missing or decreased in relative abundance in infants with CF. The consequences of these CF-specific differences in gut microbiota composition and dynamics are a delayed pattern of microbiota maturation, persistent entrenchment in a transitional developmental phase, and subsequent failure to attain an adult-like stable microbiota. We also detect the increased relative abundance of oral-derived bacteria and higher levels of fungi in CF, features that are associated with decreased gut bacterial density in inflammatory bowel diseases. Our results define key differences in the gut microbiota during ontogeny in CF and suggest the potential for directed therapies to overcome developmental delays in microbiota maturation.

## INTRODUCTION

The human gut microbiota is composed of hundreds of diverse microbial species that contribute to human health through effects on immune, metabolic, and physiological function ^1^. Alterations in the composition of the gut microbiota are linked to a variety of human diseases and it is of great basic and translational interest to understand how such altered states arise and by what mechanism they influence human health. The neonatal gut microbiota assembles at birth and proceeds to undergo dynamic and stereotypical developmental transitions through early life, defined by compositional shifts in the diversity, prevalence, and relative abundance of specific microbial taxa ^2,3^. Microbiota compositional dynamics during this period are shaped by intrinsic forces such as colonization priority effects and competition between co-resident bacteria, as well as external factors including delivery mode, breastfeeding status, and antibiotic exposure ^4^. Between three to five years of age, these dynamics stabilize into an adult-like microbiome composition ^5,6^. Disruptions to this process of gut microbiota maturation can be associated with poor health. For example, children with acute malnutrition exhibit aberrant gut microbiome maturation associated with deficits in growth as well as altered metabolic, immune, and neurological development ^7^. Correction of maturation defects via microbiota-directed dietary intervention improve child growth and development, demonstrating the interrelation between microbiota development and health ^8,9^. In mice, disruptions to microbiome maturation in early life result in heightened susceptibility to inflammation through negative impacts on immune development ^10–15^. Together, these observations provide evidence that gut microbiota maturation in early life contributes to healthy development.

Cystic fibrosis (CF) is a genetic disease defined by mutations in the gene encoding the cystic fibrosis transmembrane conductance regulator CFTR, which affects chloride ion transport across cell membranes in multiple organs ^16^. People with CF experience a range of gastrointestinal complications including chronic inflammation that resembles other intestinal inflammatory conditions and a systemic hyper-inflammatory state is present at birth, even in the absence of bacterial infections of the lung, the organ most associated with morbidity and mortality ^17–19^. Infants and young children with CF are dramatically affected and typically require intensive medical attention from birth, with nutritional and growth deficits due to pancreatic insufficiency and progressive lung disease ^20,21^. Dramatic alteration of gut microbiota composition is a hallmark of both infants and adults with CF, correlating with linear growth inhibition and increased fecal fat content and markers of intestinal inflammation ^22–28^. Notable compositional differences in the microbiota found across studies between control and CF individuals from both infant and adult cohorts include increased Proteobacteria and diminished levels of Bacteroidetes which manifest following birth and do not resolve ^22,23,26,27,29^. People with CF are often repeatedly exposed to antibiotics, pancreatic enzyme replacement therapy, proton pump inhibitor therapy, and altered diet, all factors which can plausibly alter the microbiome independently ^30,31^. However, carefully controlled gnotobiotic mouse studies in which specific pathogen free fecal material was transplanted into germfree CF or non-CF mice revealed that CFTR dysfunction alone is sufficient to drive significant differences in the microbiota ^32^. Regardless of the underlying mechanism(s), there is accumulating evidence that alterations in the gut microbiota in CF are linked to disease at distal sites ^33^. For example, reduced fecal levels of short chain fatty acid-producing gut microbes are separately associated with diminished growth and inflammation ^26,34^. Communication between the gut microbiota and the lung may also influence disease progression in CF through the systemic diffusion (or lack of diffusion) of microbially produced metabolites, or through influencing inflammation or immune cells ^33^.

The extent to which microbiota maturation defects in CF persist past 12 months is not well understood. To better define microbiota longitudinal development in CF, we performed metagenomic shotgun sequencing on fecal samples from a prospective cohort of 40 infants with CF to better define the taxonomic and functional development of the gut microbiome over the first three years of life. Here, through comparison with a large number of non-CF infant samples from previously published metagenomic datasets, we identify a delay in microbiome developmental maturation in CF that persists past 3 years of age and reveal strain-level insight into alterations in the CF gut microbiota that could be targets for therapeutic restoration.

## RESULTS

### Altered taxonomic composition of the fecal microbiota of infants with CF

We first investigated the taxonomic composition and developmental dynamics of the fecal microbiota of infants with CF. To accomplish this, we performed shotgun metagenomic sequencing on longitudinally collected fecal samples from a cohort of 40 infants with CF from northern New England from birth through the first 3 years of life. A subset of these samples were previously analyzed by 16S rRNA gene amplicon sequencing ^22,35^. We also acquired a large set of previously published metagenomics shotgun sequencing datasets derived from CF and non-CF infants comprising 3863 samples from North America and northern Europe (Supplemental Table 1) ^3,6,26, 36–38^. Taxonomic profiling confirmed previous findings that infants with CF possess dramatic phylum-level microbiota alterations compared with non-CF counterparts, with notable deficits in the abundance of Bacteroidetes and elevated levels of Proteobacteria (Figure 1A and Supplemental Figure 1, Supplemental Table 2) ^26^. Species-level tests of microbiota compositional differences in CF revealed significant separation from microbiota of non-CF infants, along a gradient of the relative abundance of *Escherichia coli*, *Ruminococcus bromii*, and *Bifidobacterium breve* (CF vs. TEDDY, unweighted UniFrac, p < 0.001, PCoA, Figure 1B) and *E. coli*, *B. breve*, and *B. bifidum* (CF vs. DIABIMMUNE, unweighted UniFrac, p < 0.001, PCoA, Figure 1C). Differences in microbiota composition between CF and non-CF samples became more pronounced with increasing age at sample collection, with older non-CF samples exhibiting greater separation (CF vs. non-CF, unweighted UniFrac, p < 0.001, PCoA, Figure 1D-E).

**Fig. 1.**
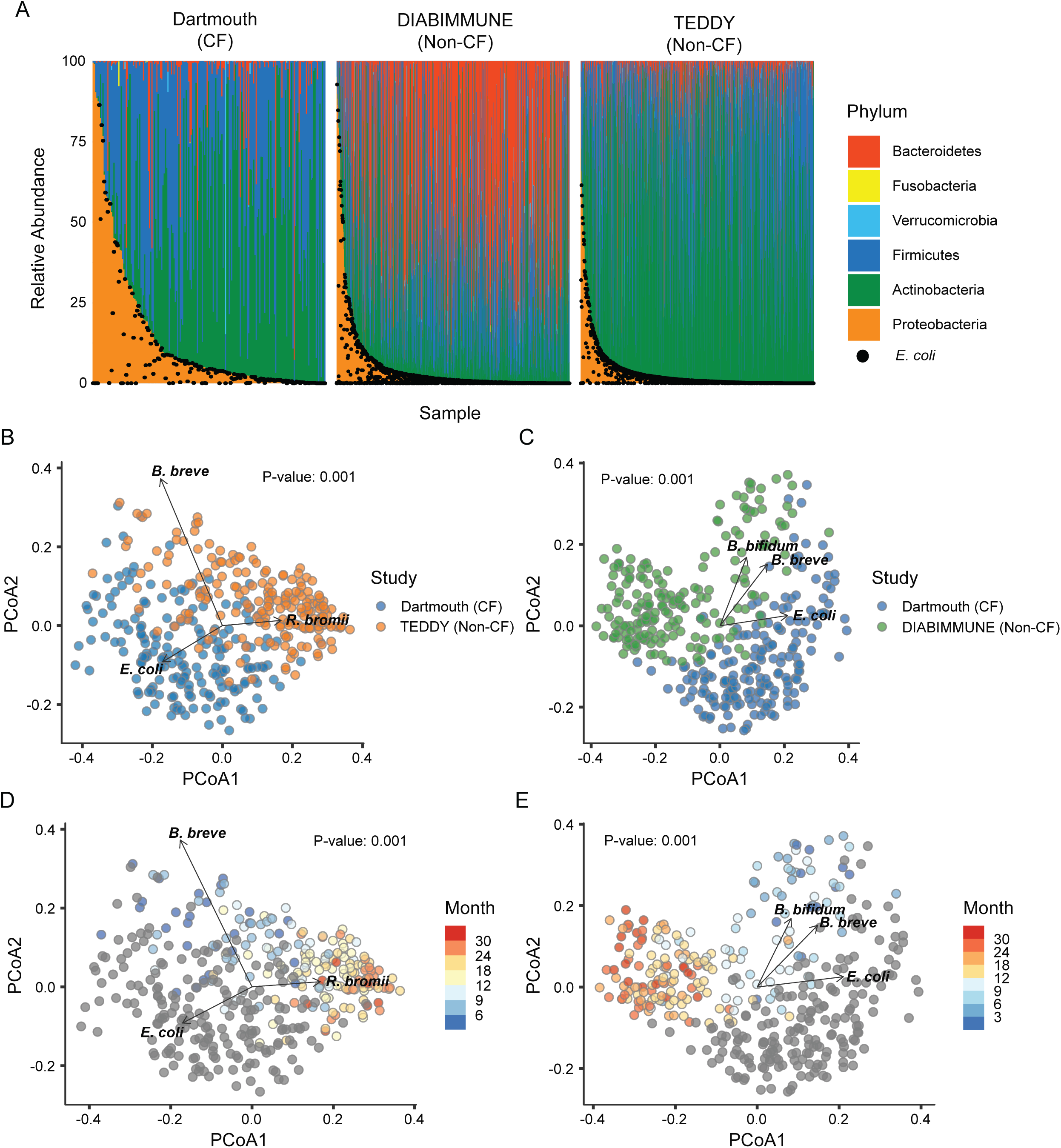
Alteration of gut microbiota composition in infants with CF. **A**, Phylum level abundance of infants in the Dartmouth (left), DIABIMMUNE (center), and TEDDY (right) cohorts from ages 0-36 months. Samples are organized by relative abundance of proteobacteria. Black dots represent the *E. coli* relative abundance for each sample (n=2589). **B-C**, PCoA of Dartmouth (blue) and TEDDY (**B**, orange) or DIABIMMUNE (**C**, green) samples from ages 0-36 months (unweighted UniFrac, PERMANOVA). A subset of non-CF samples in both cases were randomly selected to match the number of samples in the Dartmouth cohort (n = 189). Vectors indicate the top three species responsible for separation of CF samples from each of the two non-CF datasets based on Euclidean distance calculations. **D-E**, PCoA of Dartmouth and TEDDY (**D**) or DIABIMMUNE (**E**) samples from ages 0-36 months (unweighted UniFrac, PERMANOVA). Control samples were binned and colored by age with Dartmouth CF samples colored gray. A subset of non-CF samples in both cases were randomly selected to match the number of samples in the Dartmouth cohort (n = 189). Vectors indicate the top three species responsible for separation of the two datasets based on Euclidean distance calculations.

Restricting our analysis to only those samples collected up to 12 months of age, we compared microbiota composition between the two CF datasets (Dartmouth and Hayden *et al.*). We found that CF microbiomes in the first year of life share similar taxonomic profiles at the phylum level (Supplemental Figure 2A). Notably, we found that clustering of CF samples in the absence of non-CF samples revealed separation into two distinct groups defined by the presence or absence of detected *Bacteroides* (unweighted UniFrac, p < 0.001, PCoA, Supplemental Figure 2B-C). This finding is reminiscent of previously reported microbiome states observed in a non-CF infant cohort ^3^, and we similarly find that this *Bacteroides*-deficient microbiota signature was not differentiated by any measured clinical parameter including breastfeeding status, fecal fat, proton pump inhibitor usage, antibiotic treatment, or fecal calprotectin levels (data not shown).

Despite this signature of within-CF compositional divergence in the first year of life, CF infant samples from both datasets cluster separately from non-CF counterparts (unweighted UniFrac, p < 0.001, PCoA, Supplemental Figure 2D-E). Non-CF samples clustered together exhibit a high degree of overlap significance testing revealed differences largely driven by the relative abundance of Bifidobacteria (unweighted UniFrac, p < 0.001, PCoA, Supplemental Figure 2F). Together, these data demonstrate that the species-level composition of the gut microbiota of infants with CF differs dramatically from that of non-CF infants.

### Microbiome dynamics in early life

The healthy infant microbiome exhibits highly stereotypical dynamic changes in taxonomic composition over the first 3 to 5 years of life, with the most dramatic changes occurring in the first 12 months ^2,3,5,6^. Infants with CF were previously reported to possess delays in gut microbiota development in the first year of life ^26,35^. We sought to understand if such developmental delays extended past the first year of life in a different cohort. Towards this end, we first assessed family-level taxonomic profiles in the non-CF infant data and found stereotypical temporal patterns, with decreasing levels of Enterobacteriaceae over the first 12 months of age corresponding with increasing abundance of Bacteroidaceae and Lachnospiraceae (Supplemental Figure S3A). Following the first 12 months, non-CF microbiota continue to exhibit expansion of Bacteroidaceae and Ruminococcaceae populations. In contrast, this pattern was not observed for infants with CF, who instead possess high levels of Enterobacteriaceae in the first 12 months and persistently elevated levels of Bifidobacteriaceae at the expense of other families (Supplemental Figure S3A). We additionally examined changes in alpha diversity over time. In non-CF infants, as previously reported, fecal microbiome alpha-diversity increases steadily in the first year before stabilizing past 24 months of age ^5,6^. Previous reports indicated that infants with CF exhibit significantly lower alpha-diversity compared to controls ^22,26,29^. However, in contrast with these prior studies, we found no significant difference in Shannon Diversity between non-CF and CF cohorts at any time point, but significantly more detected species in CF (Supplemental Figure S3B-E, Wilcoxon ranked sum test).

### Persistent delay in microbiota maturation in infants with CF

To gain deeper insight into the nature of the altered developmental dynamics of the CF infant gut microbiota, we generated regularized random forest models as have been used previously in building models of relative microbiota age ^7,26^. The models were trained on species-level longitudinal taxonomic profiles derived from non-CF infant gut microbiome samples from the TEDDY (n=1246 and DIABIMMUNE (n=1154) studies. For each model generated from the two non-CF infant datasets, we modelled the chronological age as a function of the relative abundances of each species (546 species in TEDDY, 597 in DIABIMMUNE) at each sample collection. To evaluate model performance, we examined the correlation between predicted and true age and found that both models performed quite well (Spearman: 0.91 and 0.83, Supplemental Figure S4A-B), suggesting that there is a strong relationship between age and the developing microbiome beyond an age of 3 years. Model performance remained high even when incorporating only the top 11 or 10 most important species respectively (Spearman: 0.86 and 0.78, Supplemental Figure S4C-D, respectively). We wanted to assess whether each model contained a similar set of important species, despite being trained on completely different populations. To that end we identified the importance of age-discriminatory species for each non-CF dataset from the random forest feature importance metric and assessed correlations between the two lists. There is a high correlation between the importance scores across all species, and furthermore, 29 of the 50 most important species are shared between the two models (Spearman’s rho = 0.75, Figure 2A-B). A notorious problem with statistical models is that of extrapolation; often models perform very poorly when tested on a different population because of underlying differences in data structure between the two populations. Despite this challenge, we found that each model performed well when tested on the other population (Spearman: 0.81 and 0.75, Supplemental Figure S4E-F), suggesting that a similar set of species define microbiota age progression in different populations. In sum, we identified a set of age-discriminatory bacterial species whose prevalence and relative abundance are sufficient for accurate cross-study prediction of non-CF infant age, essentially serving as biomarkers. To our knowledge, this is the first report of a cross-study validation of a microbiota age model and our results demonstrate that such models can be highly robust and reproducible across non-CF datasets.

**Fig. 2.**
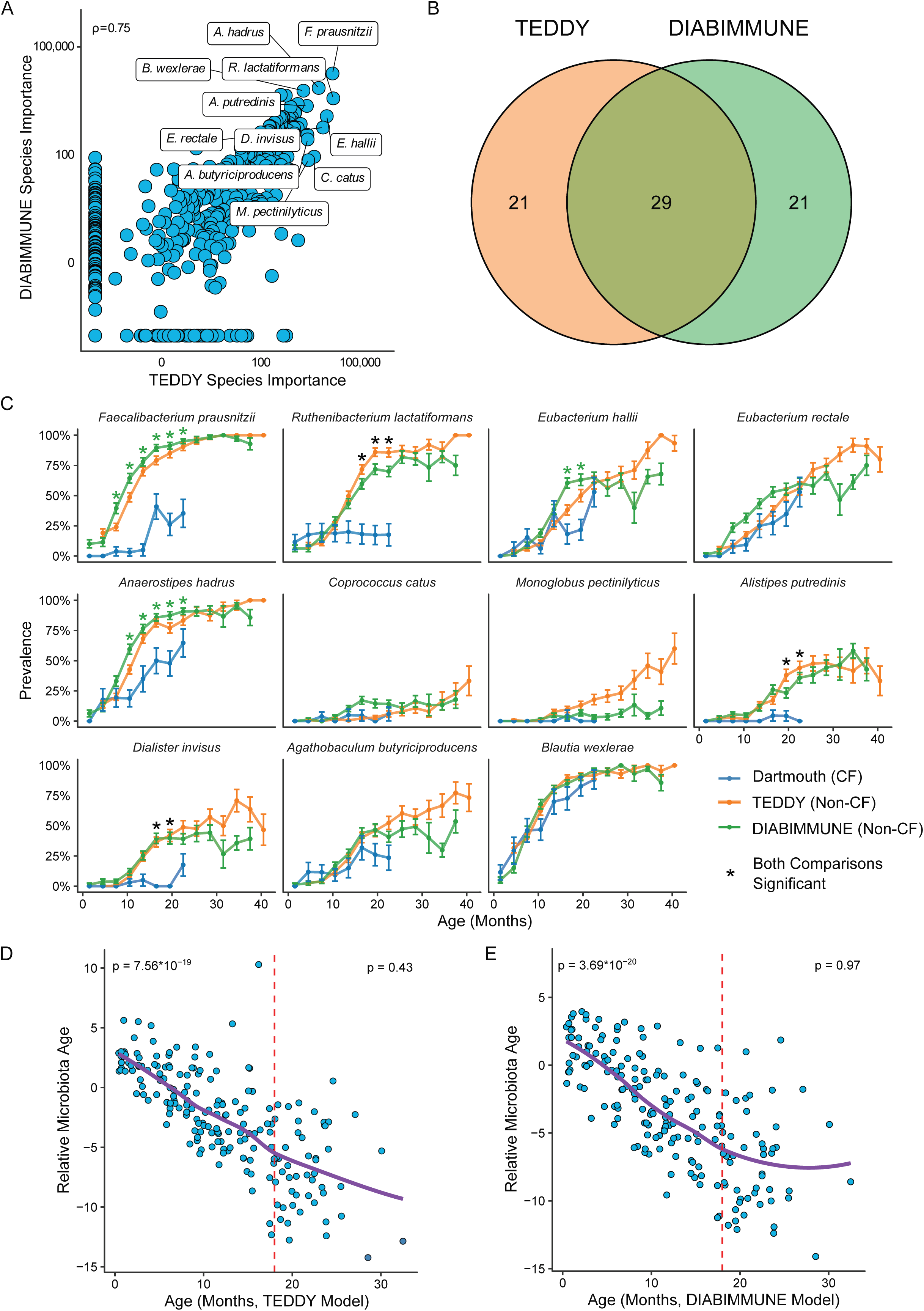
Persistent decline in microbiota relative age in infants with CF infants. **A**, Correlation plot showing species importance values from microbiota age models trained on TEDDY and DIABIMMUNE datasets. Datapoints represent individual species and the top 11 species are labeled. ρ = 0.75, spearman correlation (n = 2400). **B**, Venn diagram showing the overlap of the top 50 most important species between TEDDY and DIABIMMUNE models (n = 2400). **C**, Prevalence plots for each of the top 11 species in Dartmouth (blue), DIABIMMUNE (green), and TEDDY (orange) in the first 40 months. Stars indicate statistical significance for comparisons between Dartmouth and TEDDY (orange), Dartmouth and DIABIMMUNE (green), or both (black). *P<0.001, Fisher’s exact test (n = 2590). **D**, Relative microbiota age progression in the Dartmouth cohort when compared to the TEDDY age model. Red dotted line (18 months) signifies the time point at which the slope of the line is no longer significant. P value on upper left corner is slope prior to 18 months, P value on upper right is slope after 18 months (n = 190). **E**, Relative microbiota age progression in the Dartmouth cohort when compared to the DIABIMMUNE age model. Red dotted line (18 months) signifies the time point at which the slope of the line is no longer significant. P value on upper left corner is slope prior to 18 months, P value on upper right is slope after 18 months (n = 190).

Next, we evaluated whether age-model species were detectable in the Dartmouth CF infant samples. Several top age-model species exhibit similar prevalence between non-CF infants and infants with CF, including *Blautia wexlerae* and *Eubacterium hallii*. However, other age-model species, including the species with the highest interstudy model importance *Faecalibacterium prausnitzii*, exhibited an increase in prevalence in non-CF infants to nearly 100% by 24 months of age, yet remained at low prevalence in infants with CF (Figure 2C, Supplementary Table 3). To assess if differences in the prevalence and relative abundance of top age-model species resulted in altered relative microbiota age in infants with CF, we applied non-CF infant age models to our Dartmouth CF samples, in order to calculate the relative microbiota age for each sample, following previous methods ^7,26^. Remarkably, relative microbiota age for CF exhibited negative values that strongly negatively correlated with true age to 18 months regardless of the model used (Figure 2D-E, TEDDY age model p = 7.56×10^−19^, DIABIMMUNE p = 3.69×10^−20^). After 18 months, CF relative microbiota age remained negative but did not further decrease. These findings were consistent with age models generated for both non-CF datasets with only the top 11 or 10 age-discriminatory taxa respectively (Supplemental Figure S4G-H, TEDDY age model p = 6.14×10^−18^, DIABIMUNE p = 2.81×10^−16^), suggesting that the prevalence and relative abundance of a limited number of key taxa are biomarkers for age and can differentiate between health and disease. Together, these data indicate that the CF infant gut microbiome exhibits a persistent developmental delay in maturation that is exacerbated over time up to 18 months of age and does not recover by 36 months.

### Unsupervised clustering reveals that microbiota of infants with CF fail to transit through distinct community types

Our relative microbiota age model analysis revealed a delay in maturation of the CF infant gut microbiome compared to non-CF infants associated with a defect in the prevalence and relative abundance of key age model species such as *F. prauznitzii.* We considered two hypotheses that might explain this delay in maturation. First, the gut microbiome of infants with CF may simply be compositionally distinct from that of non-CF infants at later timepoints, with a paucity of species typically important for maturation. In support of this concept, dietary differences, disease exacerbations, and therapeutic exposures might together contribute to shift the microbiome away from a sterotypical developmental trajectory through enrichment for species such as *E. coli* ^25–27,39,40^. An alternate and non-mutually exclusive hypothesis partially supported by our analysis of beta-diversity (Figure 1D-E) is that the CF gut microbiome might initially resemble that of non-CF infants in early life but remain entrenched in an immature state at later timepoints when gut microbiome developmental maturation has continued in non-CF counterparts.

To further explore these hypotheses, and to understand in greater detail the nature of the maturation defect in CF, we employed Dirichlet multinomial mixtures (DMM) probabilistic modeling to cluster non-CF microbiomes by community types based upon the relative abundance profiles of a limited set of species identified as most important (99% of model performance, 25 species for TEDDY and 21 species for DIABIMMUNE) for both microbiota age models trained on each dataset (Figure 2A). We identified 10 discrete clusters for the TEDDY dataset and 9 clusters for DIABIMMUNE (Supplemental Figure S5). As previously reported, we found that DMM clusters generated from non-CF infant metagenomes fell into discrete phases corresponding to stereotypical progression through ontogeny, with a “developmental” phase between 0-12 months, a “transitional” phase between 12-18 months, and a “stable” phase between 18-36, respectively (Figure 3A-B, Supplemental Figure S6A-B, Supplemental Tables 4-7). Next, we assigned CF microbiome samples to clusters by using the highest probability from the non-CF model. During the developmental phase (0-12 months) we find a small skew between CF and non-CF samples in occupying the developmental clusters (67% CF vs 82% non-CF with the TEDDY model, 66% CF vs 42% non-CF with the DIABIMMUNE model). Remarkably, we found that this skew in cluster occupancy is exacerbated in CF past the transitional phase. Microbiomes from infants with CF fail to progress past clusters that are representative of the non-CF transitional phase (20% of non-CF samples in the stable phase remain in transitional clusters, compared to 61% of CF samples in the TEDDY model, 9% non-CF vs 47% CF in the DIABIMMUNE model). To investigate if this pattern of microbiome developmental progression was unique to CF, or was also observed in a separate pediatric disease, we used the same species-level taxonomic profiling and cluster assignment method to examine metagenomic samples collected from a cohort of infants prior to the onset of Celiac Disease (CD) ^41^. Unlike CF samples, those from infants with CD could be assigned to all non-CF clusters and did not display an age-dependent stall in cluster progression like that observed for CF (Supplemental Figure S7A-B, Supplemental Table 7). We conclude that our DMM cluster assignment method identifies the persistence of a transitional microbiome state during infant development that may be unique to CF.

**Fig. 3.**
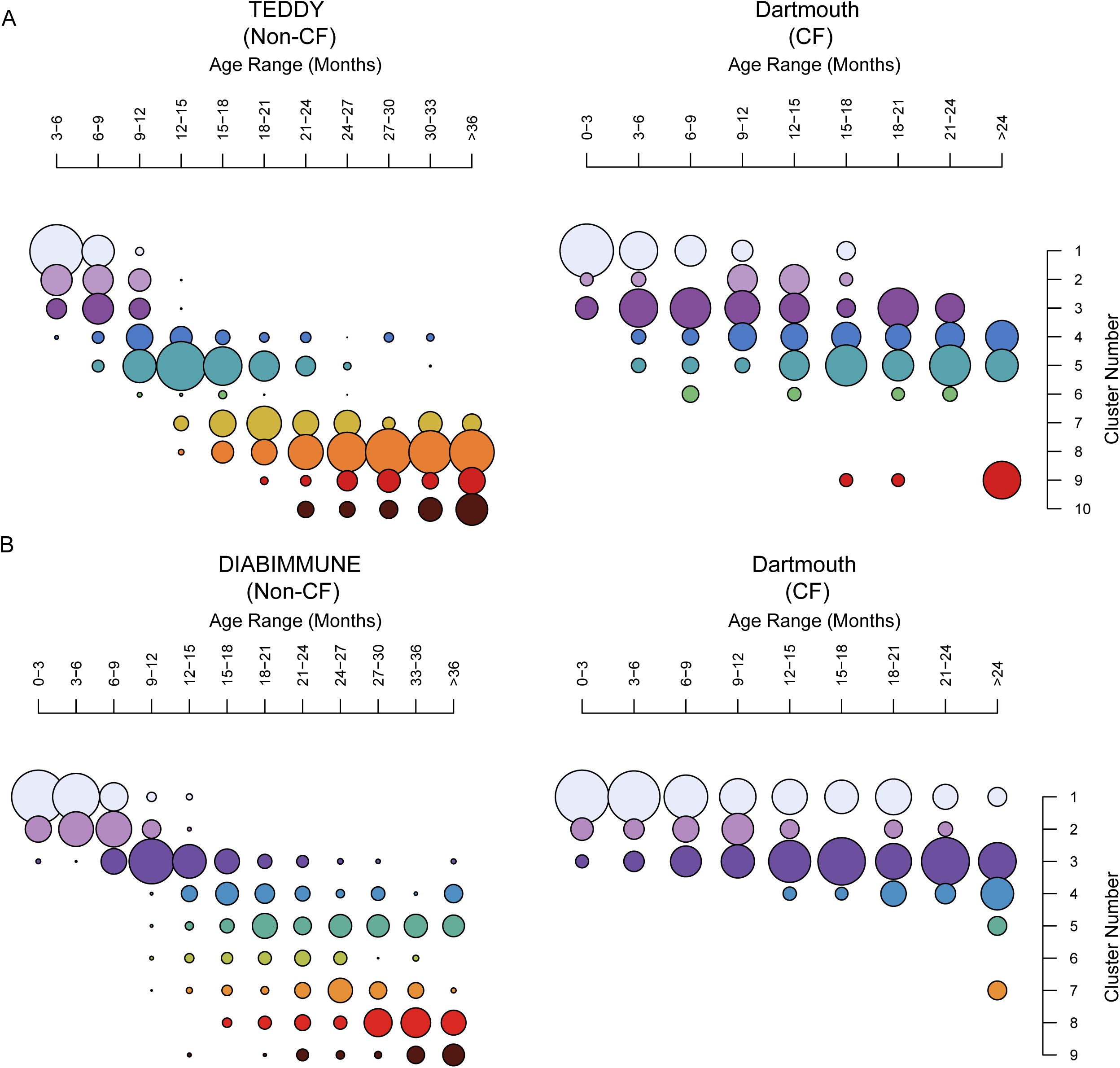
DMM clustering shows delayed development of the CF microbiome compared to non-CF controls. **A**, Dirichlet multinomial mixtures (DMMs) clustering of infants from the TEDDY (left) and Dartmouth (right) cohorts. A DMM model was trained on data from TEDDY and applied to cluster data from Dartmouth. Samples were binned into 3-month age bins and clusters were ordered based on the age bin where they are most abundant. Each cluster is denoted by a separate color (n = 1436). **B**, DMMs of infants from the DIABIMMUNE (left) and Dartmouth (right) cohorts plotted in the same way as A (n = 1344).

### Phase-level comparisons reveal a low-density microbiota signature in CF

We next assessed microbiome compositional differences by identifying species most differentially abundant in CF at each phase of development, through separate comparisons to the TEDDY and DIABIMMUNE datasets (Supplementary Tables 8-9). At each phase, species significantly depleted from infants with CF and enriched in non-CF infants included important age-model species such as *F. prausnitzii*, *Akkermansia municiphila,* and *Ruminococcus bromii* proposed to serve keystone roles in gut microbiome ecology, some of which are notable producers of host-beneficial metabolites (Figure 4A-B) ^42,43^. Additional species identified to be significantly depleted from the CF gut microbiome include prominent members of the genus *Bacteroides*, consistent with previous reports that these species are notably diminished in CF ^22,26^. Species identified to be enriched in CF across phases and in comparisons to both non-CF datasets included *E. coli*, potential pathogens like *Staphylococcus aureus* and *Clostridium perfringens*, as well as many species associated with the oral and upper respiratory microbiome. An enrichment of oral species in the fecal microbiome has been previously reported to be associated with inflammatory bowel disease and low-density gut microbiota ^44^. In line with this, we find that the sum of the relative abundance of oral species is significantly higher in CF than for non-CF infants at each phase of microbiome development, with a majority of samples exceeding an abundance threshold of 3.4% found to be associated with low-density microbiota states (Figure 5A, Supplementary Table 10) ^44^.

**Fig. 4.**
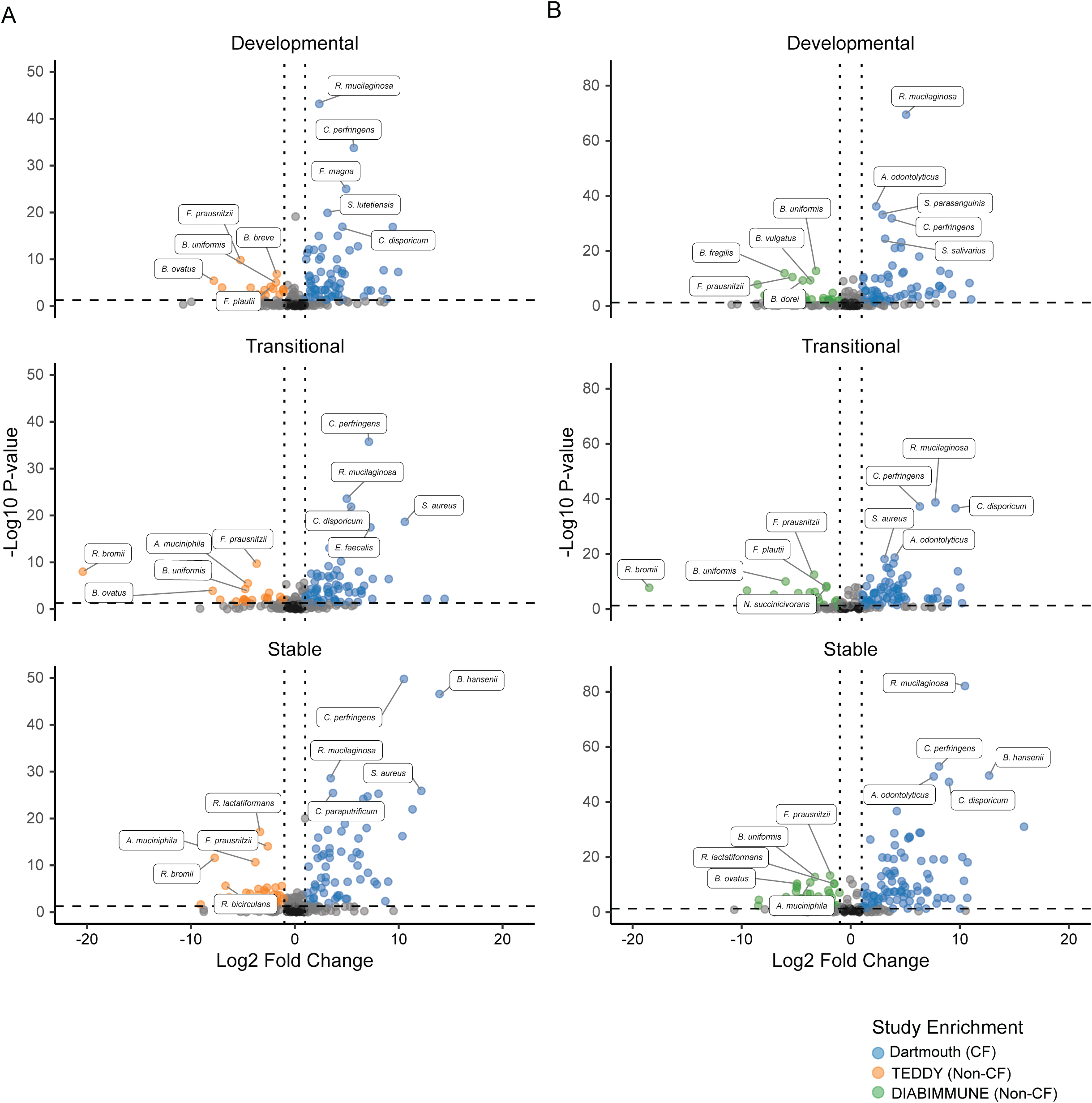
Phase-level differences in species relative abundance between CF and non-CF infants. **A**, Volcano plots of species enrichment in Dartmouth cohort compared to TEDDY cohort at each of the three developmental phases (Developmental = 0-12 months, Transitional = 13-18 months, Stable = 19-36 months). Species whose relative abundance differs significantly are colored based on which study they are enriched in (Dartmouth = blue, TEDDY = orange, P < 0.05, Wilcoxon ranked sum test, and Log2FC < −1 or > 1). The top 5 most significant species (based on P value) for each study and each phase are labelled (n=1436). **B**, Volcano plots of species importance in Dartmouth cohort compared to DIABIMMUNE cohort at each of the three developmental phases (Developmental = 0-12 months, Transitional = 13-18 months, Stable = 19-36 months). Species of statistical significance (P < 0.05, Wilcoxon ranked sum test, and Log2FC < −1 or > 1) are colored based on which study they are enriched in (Dartmouth = blue, DIABIMMUNE = green). The top 5 most significant species (based on P value) for each study and each phase are labelled (n=1344).

**Fig. 5.**
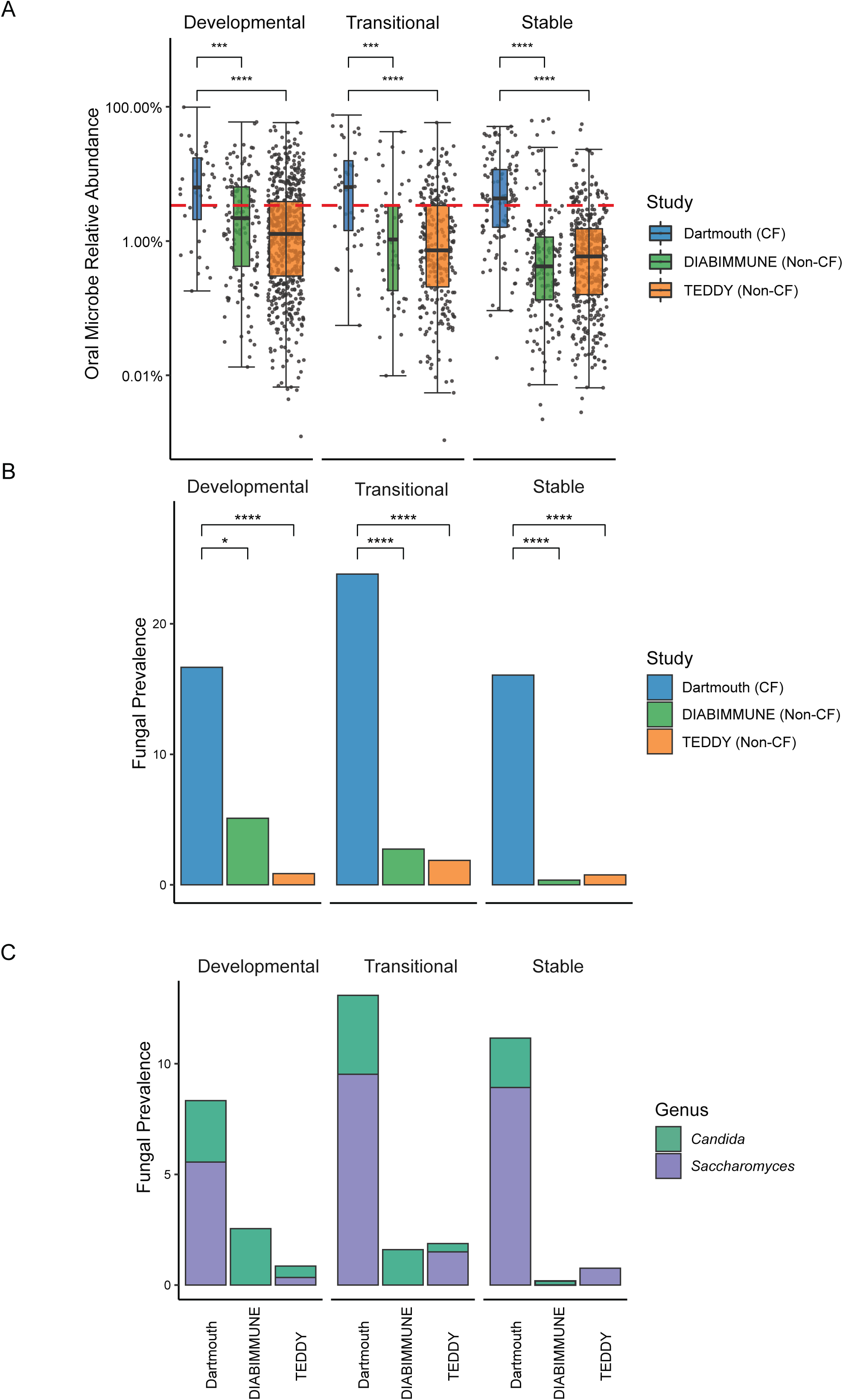
Oral and fungal relative abundance is higher in CF infants compared to non-CF infants in early life. **A**, Relative abundance of oral microbes in Dartmouth (blue), DIABIMMUNE (green), and TEDDY (orange) cohorts at each developmental phase (Developmental = 0-12 months, Transitional = 13-18 months, Stable = 19-36 months, n=2590). Red line signifies 3.4% relative abundance. **B**, Fungal prevalence in the Dartmouth (blue), DIABIMMUNE (green), and TEDDY (orange) cohorts at each developmental phase (Developmental = 0-12 months, Transitional = 13-18 months, Stable = 19-36 months). Abundance was first collapsed by genus and prevalence was calculated per individual. **C**, Fungal prevalence by genus in the Dartmouth, DIABIMMUNE, and TEDDY cohorts at each developmental phase (Developmental = 0-12 months, Transitional = 13-18 months, Stable = 19-36 months).

An increase in fungi in the gut has also been reported for microbiomes with low bacterial density. We assessed fungal prevalence in our datasets and found elevated fungi in CF compared to either non-CF dataset, with frequent detection of *Saccharomyces* and *Candida*, both of which have independently been associated with inflammatory bowel conditions (Figure 5B-C, Supplementary Table 11-12) ^45,46^. Together, these findings implicate a low-density gut microbiota in CF, with corresponding loss of beneficial species that include prominent contributors to non-CF microbiota relative age.

### Altered microbiota functional capacity in CF

Differences in the composition of the microbiota can result in altered functional potential between groups. To assess if this is the case, we next evaluated whether alterations in the fecal microbiota of infants with CF resulted in significant differences in metabolic potential and functional pathway abundances through comparisons to both TEDDY and DIABIMMUNE (Supplemental Figures S8–9, Supplemental Tables 13-14). We used hierarchical clustering of the top most-significant functional modules from comparisons of CF to each non-CF dataset separately. This analysis clearly showed separation by health status instead of sample age, with the exception of the CF stable phase which clustered together with the TEDDY transitional and stable phase samples. Pathways determined to be the most significantly enriched in CF at all phases include fatty acid biosynthesis and the dicarboxylate-hydroxybutyrate cycle and reductive acetyl-CoA (Wood-Ljundahl) pathways, indicating a dramatic shift in metabolic capacity in CF previously associated in non-CF infants with the pre-weaning state ^2,47,48^. In contrast, modules involved in the adult microbiome-associated vitamin B7 biotin biosynthesis pathway were significantly depleted in CF samples across phases ^48^. We also assessed the functional contribution of taxa determined to be statistically enriched in either non-CF or CF infants for each phase of microbiome development. Notably, species enriched in healthy infants were major contributors to functional pathway abundance (Supplemental Figure S10A-B), mirroring the high relative abundances of these species (Supplemental Figure 10C-D). *F. prausnitzii* was particularly notable as a top functional pathway contributor across phases and in both non-CF dataset comparisons. In contrast, species enriched in CF exhibited minimal contributions to functional pathway abundance and were on average low abundance (Supplemental Figure S10A-D). Together, these findings demonstrate that the alterations in taxonomic composition we observe between non-CF infants and infants with CF reflect dramatic differences in functional pathway abundance.

### Reduced strain-level diversity of *Faecalibacterium prausnitzii* in CF

We chose to investigate the CF-depleted butyrate producer *Faecalibacterium prausnitzii* in greater detail at the strain level, due to its importance in our microbiome age-model analysis (Figure 2), its significant contribution to functional pathway abundance in non-CF datasets (Supplemental Figure S10), and its well-established role in gut health and development ^42^. *F. prausnitzii*, has previously been identified as a biomarker of intestinal health and may limit inflammation in part through production of the short chain fatty acid butyrate ^49^. Healthy infants are typically colonized early in life with *F. prausnitzii*, with the relative abundance and strain diversity increasing over age ^50^. We therefore assessed *F. prausnitzii* diversity in infants with CF, comparing to the TEDDY and DIABIMMUNE non-CF cohorts. We detected all human-associated *F. prausnitzii* clades in at least one infant metagenome, with the prevalence of clades B, C, D, G, and I increasing with age (Supplemental Figure S11A-B). Comparing *F. prausnitzii* diversity across samples, we found all clades to be more prevalent in the non-CF cohorts except for clade L, which is more prevalent in both distinct CF cohorts (Figure 6A-B). Interestingly, clade L is predominantly associated with pets including canines and felines ^50,51^ and pet-associated *F. prausnitzii* has been reported to colonize humans ^52,53^. Phylogenetic analysis of *F. prausnitzii* clade L strains including those detected in CF infant samples, reveals relatedness between human- and pet-derived strains (Figure 6C). These data suggest the CF infant gut is more permissive to colonization by non-human strains of *F. prausnitzii* than that of non-CF infants. Finally, since decreased within-sample clade diversity of *F. prausnitzii* is associated with disease states including inflammatory bowel disease ^50^, we compared *F. prausnitzii* diversity between non-CF and CF samples for each phase. We detect significantly more clades in non-CF samples than in CF for all phases (Figure 6D). Together, these results demonstrate that a prominent feature of the CF infant microbiome is the dramatically altered prevalence and diversity of *F. prausnitzii*.

**Fig. 6.**
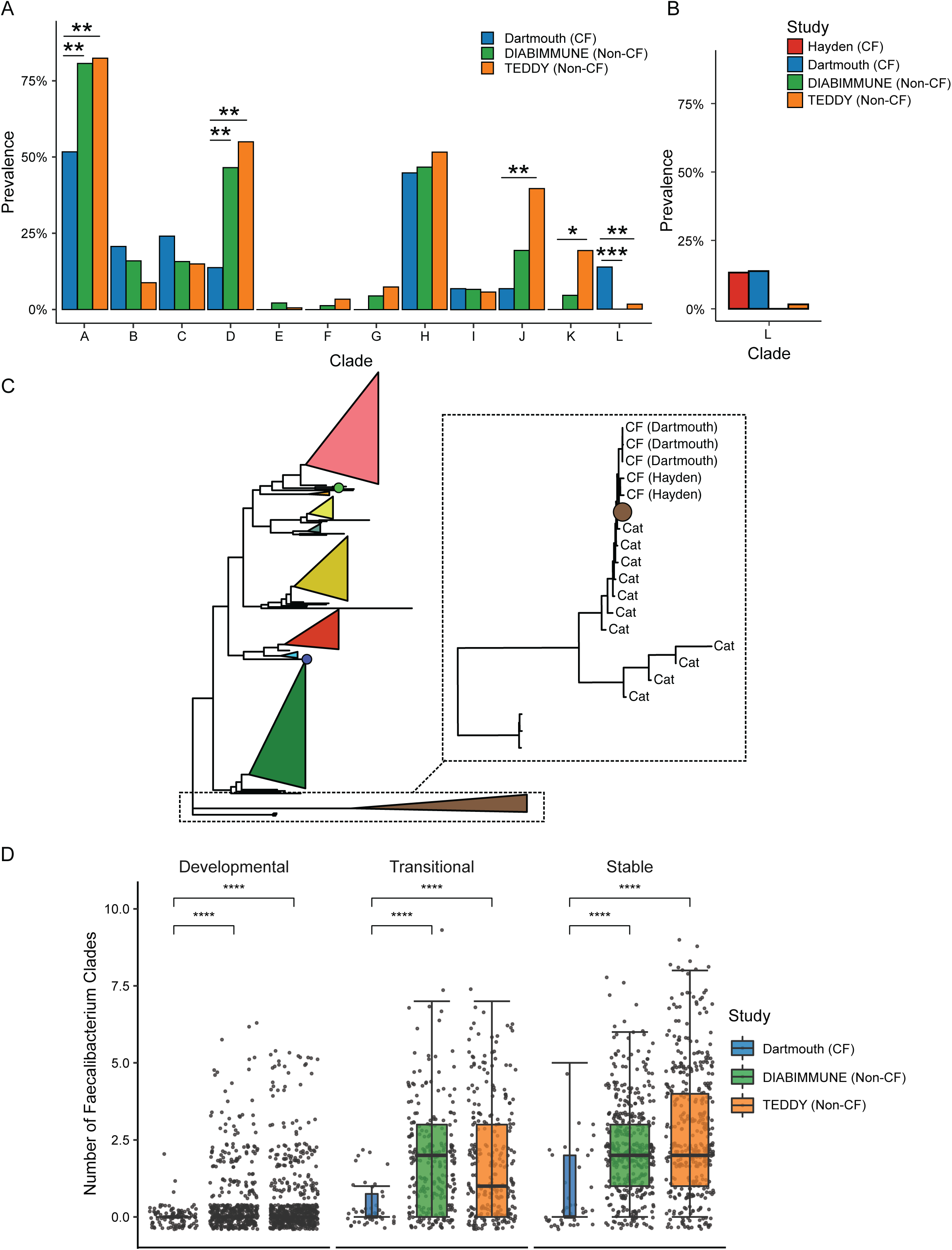
*Faecalibacterium prausnitzii* strain diversity is altered in CF infants compared to non-CF controls. **A**, Overall prevalence of *F. prausnitzii* clades in CF and non-CF cohorts (*FDR <= 0.05, Fisher’s exact test). **B,** Clade L prevalence across CF and non-CF cohorts. **C**, Phylogenetic tree of *F. prausnitzii* strains. The tree contains data derived from publicly available reference genomes as well as sequences reconstructed from feline metagenomes and human CF infant metagenomes (Dartmouth and Hayden). Detected strains were assigned to clades based on the presence of clade specific marker genes and collapsed into triangle form. Black box highlights strains in clade L and is magnified in inset box to the right depicting strains inferred from feline metagenomes. **C**, Number of *F. prausnitzii* clades identified in Dartmouth (blue), DIABIMMUNE (green), and TEDDY (orange) cohorts at each developmental phase (Developmental = 0-12 months, Transitional = 13-18 months, Stable = 19-36 months). ****P<=0.0001, Wilcoxon ranked sum test (n=2590).

## DISCUSSION

Our findings demonstrate that the gut microbiota of infants with CF fails to undergo typical developmental maturation, instead remaining entrenched in a transitional-like community state. This conclusion is grounded upon our robust age-model analysis benchmarking non-CF infant microbiome development across two of the largest published publicly available infant metagenomic datasets, TEDDY and DIABIMMUNE. Cross-validation of age-model species importance demonstrates that key taxa are reproducibly associated with chronological age across non-CF datasets that are compositionally distinct with major differences in prominent phyla such as Bacteroidetes and Bifidobacteria. While the prevalence and relative abundance of species like *F. prausnitzii* therefore represents a biomarker of age, it is important to note that many important age-model species are themselves found at high levels in the adult microbiome and have been implicated in vital functions such as the production of SCFA and other metabolites that impact host physiology at both local and distal sites ^54,55^. Altered microbiome maturation therefore is likely to reflect fundamental consequences for both taxonomic composition and function.

The gut microbiota are particularly critical for early-life training of the developing immune system and disruptions to the microbiota during this time period can lead to subsequent metabolic and immunologic dysregulation ^10,56^. In mice, microbiome dynamics during the weaning period are necessary for a transient inflammatory response known as the weaning reaction. Suppression of microbiome dynamics (specifically the bloom in Gram-positive SCFA producers during and following weaning) prevents the weaning reaction and predisposes mice inflammation and pathogen susceptibility via restriction of peripheral regulatory T cell expansion ^10,13^. We find that infants with CF harbor populations of Gram positive SCFA producers like *F. prausnitzii* fail to expand in relative abundance over the first three years of infancy and consequently do not diversify at the strain level as in non-CF infants. The potential for a link between impaired microbiome development in CF and deleterious immunological sequelae warrant further investigation.

We identify an enrichment of oral-derived bacterial taxa at all developmental phases in infants with CF. This signature has been implicated in a low-density gut microbiota state ^44^. In mice, experimental manipulation of gut microbiota density results in changes to host metabolism and immunological dysregulation ^57^. One potential consequence of reduced gut microbiota density is lowered concentrations of microbiota dependent metabolites that mediate beneficial effects for the host. Altered metabolite concentrations have been observed in infants with CF, however, if these changes are due to alterations in taxonomic composition, altered density, or both remains an open question ^58^. Another consequence of low gut microbiota density is an increase in the abundance of fungi, and we observe increased prevalence of fungal taxa such as *Candida* in infants with CF, consistent with evidence that people with CF are more likely than healthy counterparts to possess anti-*Saccharomyces cerevisae* antibodies (ASCA), which recognize oligomannose in the yeast cell wall, including that of *Candida* species ^59^. Intestinal colonization by *Candida* species is linked to inflammatory bowel diseases and the induction of Th17 responses both in the gut and in the lung, however the impact of infant carriage of Candida in CF is unexplored ^60^.

Our study does not incorporate clinical metadata such as *CFTR* genotype, pancreatic insufficiency status, fecal fat, inflammation, or linear growth metrics. While these factors may contribute to or are known to be associated with microbiome composition in some cases, our study is not sufficiently powered to assess their independent relative contributions to the patterns we observe. It is therefore striking that despite ignoring clinical parameters, we detect dramatic differences in microbiome taxonomic composition and functional potential along with altered microbiome developmental dynamics in infants with CF compared to non-CF counterparts. Notably, exposure to antibiotics has been previously associated with a similar magnitude of compositional effect in CF as in control infants, suggesting that antibiotic use alone was not likely to contribute to the differences that we detect ^25^. Together with experimental studies in mice demonstrating an association of *CFTR* genotype altered microbiome composition ^32^, we propose that our data support that impaired CFTR function alone may be sufficient to drive at least some aspects of altered microbiota maturation. Further investigation using animal models of CF will be needed to understand the causal underpinnings of CF-related microbiota alterations and their consequences for the host more fully.

The link between atypical microbiome maturation and intestinal inflammatory diseases in infants remains generally underexplored. Our analysis of microbiome development in infants prior to onset of Celiac Disease (CD) revealed no obvious differences compared to non-CF development. While it is possible that impaired microbiome development is a unique feature of CF, an alternate possibility could be that delays in microbiome development reflect active disease during this time-period. In CF, infants exhibit levels of fecal calprotectin that increase over time and other markers of inflammation from very early in life and it is probably that active disease in CF initiates at or before birth ^61^. Although CD and other inflammatory intestinal diseases typically manifest later in childhood at the earliest, it is possible that disease occurring during the developmental or transitional phase of development (rather than after the attainment of stable phase) might be associated with patterns maturation delay similar to what we observe in CF. It is notable that the infant disease kwashiorkor, unrelated to CF and caused by severe malnutrition, is also linked to maturation delays in microbiome development and coincident intestinal inflammation ^62^. Future mechanistic work investigating the link between microbiota age delay and disease in CF will benefit from the experimental framework leveraging gnotobiotic models and dietary interventions laid out by the kwashiorkor field ^63^. Nutritional supplementation in CF targeted at promoting the growth of keystone age-model species like *F. prausnitzii* may be a promising avenue to overcome microbiome age delay in hopes of improving health.

## METHODS

### CF cohort sample collection

This study was approved by the Dartmouth College Committee for the Protection of Human Subjects (CPHS Study # 00021761). Infant fecal samples were collected longitudinally from 13 days up to 45 months of age. A total of 190 samples were analyzed. Each subject had between 1-11 samples collected, with a median of 5 samples collected per subject. Fecal samples were collected by parents and initially stored in a home freezer prior to transfer to clinics throughout New England during routine visits with clinicians. At the clinic, the samples were stored at −80°C until they were transported to the lab to be aliquoted, stored in −80°C and later processed for sequencing.

### Non-CF control cohort selection

Longitudinal control studies were selected based on the sample size and age range of individuals. We specifically selected cohorts ranging between 0 and 3+ years of life, and who had large sample sizes of infants that had at least two samples collected over that time frame. Hayden et al. was selected to compare CF cohorts as well as our CF cohort to an age matched healthy cohort within the first year of life ^26^. All samples from the DIABIMMUNE study ^3,36,38^ and 10% of the individuals followed in the TEDDY study ^6,37^ were selected as large healthy cohorts extending into the third year of life as an extended age matched control to our cohort. To obtain a representative number of samples from the TEDDY study, 88 individuals were randomly selected and evenly distributed from Europe and the United States to account for any possible location bias. All collected samples from each individual were used in subsequent computational analysis.

### DNA isolation and sequencing

Infant fecal samples stored at −80°C was thawed and aliquoted to ~100mg in Eppendorf tubes prior to DNA extraction. DNA was isolated using the Zymo Quick-DNA Fecal/Soil Microbe Miniprep Kit (Cat #D6010). DNA concentrations were measured via Qubit fluorometer and sequencing performed on an Illumina NextSeq platform (Microbial Genome Sequencing Center (MiGS), now SeqCenter). Sequencing produced an average of 20,320,837 reads and 97.8% of samples had at least 12 million reads.

### Taxonomic profiling

After sequencing, samples were filtered and trimmed when applicable using BBDuk, trimming bases with a quality score below 12 and filtering out reads below 20 bases. BBMap was used to remove human DNA from each sample prior to taxonomic profiling ^64^. Quantification and profiling of taxa in each sample was performed using MetaPhlAn3 ^65^. Briefly, MetaPhlAn3 identifies members of the microbiota in each sample using a database comprised of unique marker genes specific to each individual species, which are used to quantify the abundance of each microbe detected in sequencing ^66^. We used default settings and a minimum read length of 30. For the phylum-level plots in Figure 1, samples were collapsed and summed by phylum classification. After filtering for top phyla, samples were normalized to 100% and were rank ordered by the abundance of Proteobacteria, with *E. coli* abundances determined for each sample prior to plotting. Raw taxonomic relative abundances were used for all subsequent analyses. The DMM clustering method (see below) required counts, and we estimated the number of counts mapping to each lineage with the rel_ab_w_read_stats flag.

### Statistical analysis

Statistics were performed using R version 3.6.3 and version 4.1.2 (R Core Team 2021). Significance was considered at P < 0.05 unless otherwise indicated. Multivariate analysis was performed using PERMANOVA tests implemented by adonis in vegan version 2.5.7 ^67^. Multidimensional scaling (principal coordinate analysis) of taxonomic profiles was completed using unweighted Unifrac in phyloseq version 1.38.0 and the top three species were identified by calculating the mean euclidean distance of the vectors for each species ^68^. Alpha diversity was calculated using Shannon diversity index using custom code. Hypothesis testing was performed using either a one-sided Wilcoxon rank-sum test or Fisher’s exact test, as indicated. Multiple hypothesis correction was performed using the Benjamini-Hochberg procedure where applicable. To identify significant associations between KEGG modules and developmental phases, we used MaAsLin2 for linear mixed-effects modeling, using module and phase as fixed effects and a minimum abundance of 0.01%, and the significant-results file from the automated output was used for further analysis ^69^.

### Relative microbiota age analysis

We implemented the age model as has been described previously ^7,26^. In brief, we used regularized random forests (RRF package in R) to predict the age of a microbiome sample from the relative abundance of microbiome species (calculated by MetaPhlAn3). Conceptually, the relative microbial age is the residual of the model prediction, but there is an additional step to correct for the significant saturating effect in the predicted model age. Specifically, we used a third-degree spline to model the relationship between the true age and the predicted age, then we took the difference between the predicted age from the random forest and the prediction from the spline. The importance of each species was calculated using the random forest approach to variable importance (the decrease in residual sum of squares when splitting on the variable, averaged over all trees). We identified a subset of species that gave either 95% of maximum performance (10 or 11 species for TEDDY and DIABIMMUNE, respectively) or 99% of maximum performance (25 species) by step wise addition of species in order of decreasing importance until we reached the performance cutoff (either 95% or 99%). The performance of the model was evaluated using the spearman correlation between the predicted and actual age of the sample. All performance was evaluated in 10-fold cross validation within each respective dataset.

### Dirichlet multinomial mixture modeling

We followed previous methods to cluster developing infant gut microbiome data using the Dirichlet Multinomial Mixtures (DMM) approach as implemented in the DirichletMultinomial package in R ^6,70^. We provided the clustering algorithm with a subset of species to focus on the most important species for development. Specifically, we only considered the limited set of species that produced 99% performance in the age model (see above). Alternative models trained with either the top 25 most abundant species, or all species with more than 0.01% average abundance gave qualitatively similar results (data not shown). DMM is a count-based method and we used the count data estimated by MetaPhlAn3 (see above). We identified the optimal number of clusters using Bayesian Information Criterion (BIC), however we found that the optimal cluster number was unstable with +/- 3 clusters changing over replicate runs. In each 3-month bin of data, we calculated the fraction of samples in each cluster. Bins with less than 15 samples were dropped due to statistical issues arising with low sample sizes. Clusters were ordered by the time point bin at which they were most abundant. We plotted the percentage of samples in each age bin as circles using the igraph package in R. Following previous methods, we dropped low abundance assignments (less than 4%) ^6^, but contrary to their work we did not plot temporal transitions between clusters because we did not have sufficient sample density from each CF individual.

### Oral and fungal microbiota analysis

To calculate the oral bacterial fraction of metagenomic samples, we filtered taxa by the genera listed in Liao *et al*. and summed them on a per sample basis ^44^. Samples were then filtered for those containing greater than 3.4% oral species to find the percentage of total samples with a significant fraction of oral microbes (Supplementary Table 9).

Fungal prevalence was calculated by isolating all Eukaryota data from the MetaPhlAn3 and finding the number of samples for which the fungal abundance was > 0 and dividing by the total number of samples, followed by converting into a percentage. This was done after summing the genera on a per sample basis for a total fungal prevalence calculation and at the individual genus level. Statistical tests comparing prevalence between healthy and CF datasets were performed using Fisher’s exact test.

### Functional Pathway Analysis

The functional capacity of each sample was analyzed using HUMAnN3, which aligns reads to a database of genes belonging to members of the microbiota ^71^. Briefly, identified genes are quantified at both the community and individual level, which we then annotated using the Kyoto Encyclopedia of Genes and Genomes (KEGG). Community level values were then collapsed into functional pathways and modules for further analysis. Statistical analysis was performed using MaAsLin2 using default parameters, with a minimum abundance of 0.01% ^69^. In short, MaAsLin2 uses linear modeling to account for covariates that may impact the desired result. Disease state and age were accounted for in this process and were used as fixed effects. Heatmaps were generated using only modules noted in the significant results file output, and module abundances were rescaled to 1 to show contrasts on a per module basis.

To select the most significant species for the functional bar and box plots, significance testing was run at the species level using the taxonomic abundances generated by MetaPhlAn3 using the healthy cohort as the control population. Significance was defined by having a P value < 0.05, and a Log2FC of greater than 1 or less than −1. These values were then plotted in volcano plots to demonstrate species enrichment or depletion in the CF cohort. The top five most significant species in both the healthy and CF datasets for each comparison and each developmental phase were labeled on the volcano plots and were selected for further analysis. These species were filtered from the HUMAnN3 analysis, and this data was collapsed into KEGG pathways by finding the mean abundance among all samples where the functional abundance was > 0, then summing the abundances for each module within each pathway for plotting. Prevalence was calculated by summing the total number of species with relative abundance > 0 divided by the total number of samples and multiplying by 100 to obtain percentage values.

### Strain-level analysis of Faecalibacterium prausnitzii

To detect *F. prausnitzii* strains we used a set of clade-specific marker genes that were previously defined ^50^. We aligned metagenomic reads to the marker genes using bowtie2 (2.3.4.1, with parameters -a -N 1) and quantified the number of reads mapping to each maker, discarding multi mapping reads, reads that mapped at less than 95% identity, 50% of the read length, or below a quality score of 20. We determined the presence of a marker gene if it had at least 25 mapped reads, and we determined the presence of a strain if we could identify at least 10% of the marker genes. This is in contrast to previous methods that used the same cutoff of 10% of the marker genes to determine the presence of a strain, but they determined gene presence using a consensus sequence-based approach, specifically a consensus sequence with at least 95% identity over 50% of the gene length. We experimented with both approaches and found cases where divergent strains hovering around the 95% identity threshold would be missed by the consensus approach but successfully found by the count-based approach, and that these cases were convincing because of a large number of read counts. Additionally, the number of clades were quantified using the marker gene approach in each dataset at each developmental phase. Statistical comparisons between the prevalence of clade were performed with the chi squared test for the developmental stages, or the fishers exact test for the difference between CF and healthy. Differences in the number of strains was assessed using a one-sided Wilcoxon ranked sum test.

We built a phylogenetic tree of *Faecalibacterium prausnitzii* by obtaining 367 reference genomes from Genbank in September 2022. We used prokka (1.13) to infer gene content and roary (3.13) to infer core genes ^72,73^. For phylogenetic tree construction we used core genes that were present in at least 50% of the genomes. To construct the tree, we identified marker genes in genomes using BLASTN against the prokka inferred gene sequences and filtered hits with less than 90% sequence identity and covering less than 90% of the length of the query and the hit. In addition, we included sequences inferred from CF-derived metagenome datasets, as well as previously sequenced cat metagenomes ^51^. Marker genes in metagenomes were inferred by aligning reads to the marker sequences using bowtie 2, then constructing pileups using samtools (1.9) and creating a consensus sequence with the major allele at every position along the gene sequence ^74,75^. Positions below a coverage of 5, or where the major allele was less than 5 counts higher than the minor allele were replaced with an N, as well as positions 10 bases from the start and from the end of the gene. We aligned all representatives of a marker gene individually using MAFFT (v7.505) and concatenated the individual alignments together. Using the concatenated alignment we created a phylogenetic tree using RAxML using the GTRGAMMAI (general time reversible model with a gamma model of rate heterogeneity and with an estimate of the proportion of invariable sites) ^76^. Additionally, the number of clades were plotted in each dataset at each developmental phase on a per sample basis. Statistics were performed using a one-sided Wilcoxon ranked sum test.

## Data Availability

The shotgun metagenomic samples generated in this study are publicly available through the Sequence Read Archive at the National Center for Biotechnology Information (NCBI) under BioProject accession number PRJNA955235. Other data analyzed in this study are listed in Supplemental Table 1.

## Code Availability

Detailed analysis scripts are available upon request.

## Author contributions

Conceptualization: BDR, PES, AJV. Methodology: PES, AJV. Investigation: PES, AJV, CEP, RAV, KAB. Visualization: PES, AJV. Supervision: BDR. Writing—original draft: BDR, PES, AJV. Writing—review & editing: BDR, PES, AJV, CEP, RAV, KAB, GOT, JCM.

## Competing interests

No competing interests are declared by the authors.

## Acknowledgements

This project was supported by generous funding from the Cystic Fibrosis Foundation (ROSS20R3, OTOOLE19GO, MADAN18GO, MADAN18AO, 00389A122MADAN, STANTO19R0) and National Institutes of Health (P30-DK117469, T32-AI007363, T32-HL134598, UH3OD023275) as well as startup funding from the Dartmouth College Geisel School of Medicine. We gratefully thank the participants and families of the studies. We are also grateful for helpful advice and constructive feedback from Anne Hoen, Christopher Stewart, Zachary Sellers, and members of the Dartmouth College DartCF, M2P2, and JEMM communities. This research utilizes data obtained by the TEDDY study group, a collaborative clinical study sponsored by the National Institute of Diabetes and Digestive and Kidney Diseases (NIDDK), National Institute of Allergy and Infectious Diseases (NIAID), National Institute of Child Health and Human Development (NICHD), National Institute of Environmental Health Sciences (NIEHS), and Centers for Disease Control and Prevention (CDC), and JDRF. The data from the TEDDY study reported here were supplied by the database of Genotypes and Phenotypes (dbGaP), which is maintained by the National Center for Biotechnology Information (NCBI). This manuscript was not prepared in collaboration with investigators of the TEDDY study and does not necessarily reflect the opinions or views of the TEDDY study, dbGaP, or the NIDDK.

---

**Fig. S1.**
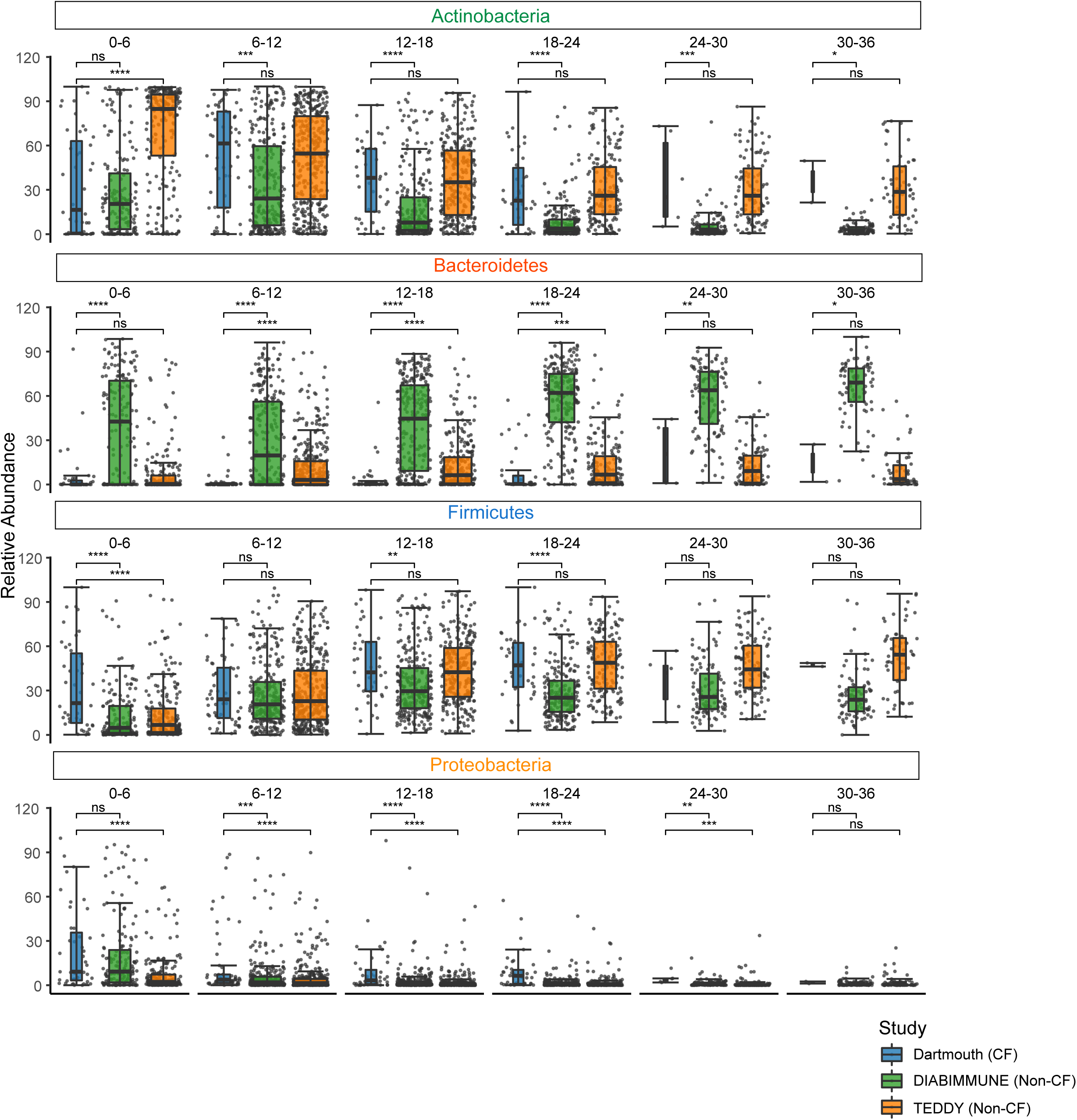
Relative abundance of select phyla differ between infants with CF and non-CF controls. Select phylum level relative abundance from infants in the Dartmouth (blue), DIABIMMUNE (green), and TEDDY (orange) cohorts over time. Samples were placed into six-month age bins and are shown as boxplots. *P <= 0.05, **P <= 0.01, ***P <= 0.001, ****P <= 0.0001, Wilcoxon ranked sum test (n=2589).

**Fig. S2.**
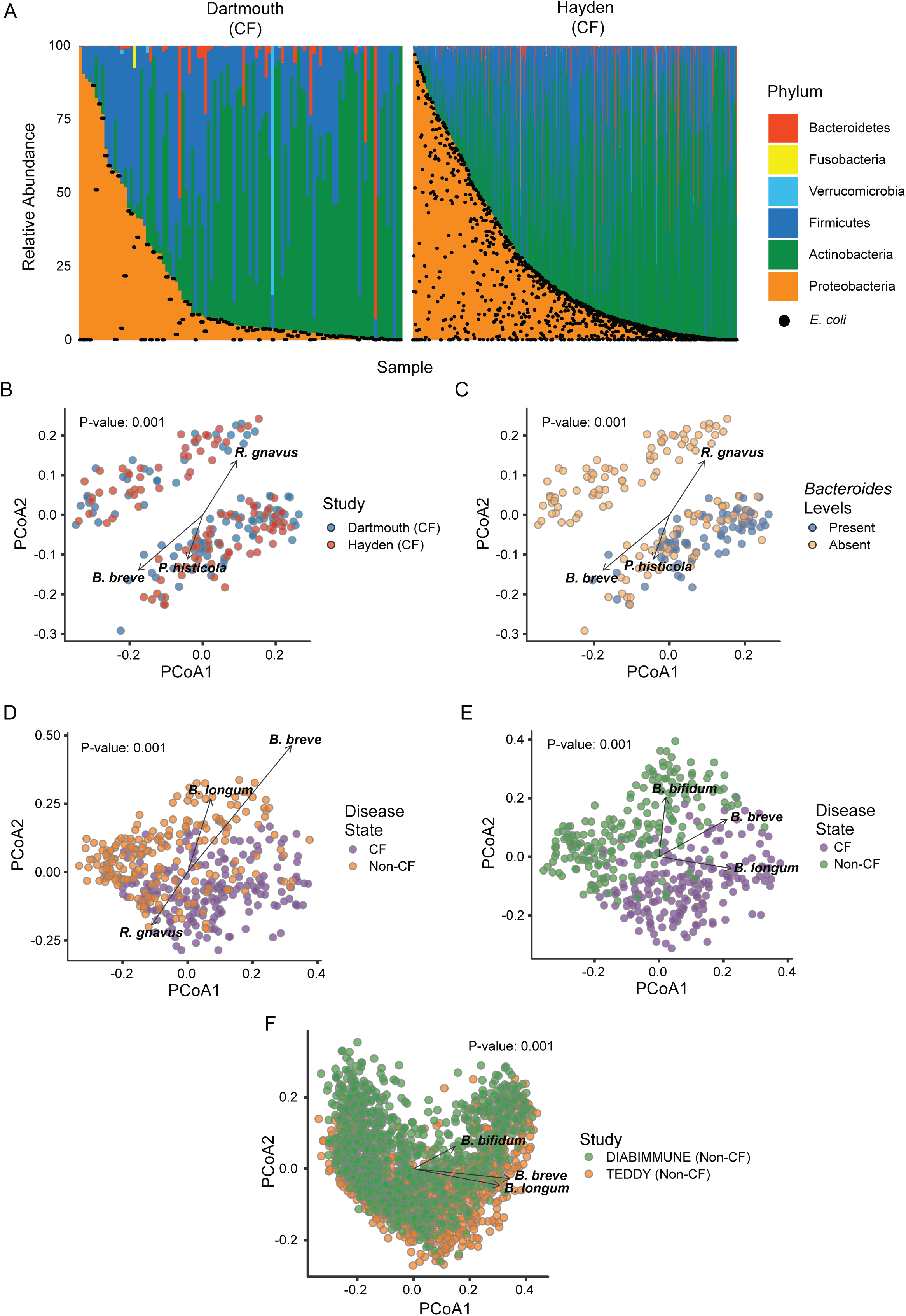
Gut microbiota from CF cohorts exhibit compositional similarity in the first year of life. **A**, Phylum level relative abundance of infants in the Dartmouth (left), Hayden CF (right) cohorts from ages 0-12 months. Samples are organized by relative abundance of Proteobacteria. Black dots represent the *E. coli* relative abundance for each sample (n=1255). **B**, PCoA of Dartmouth (blue) and Hayden CF (red) samples from ages 0-12 months (unweighted Unifrac, PERMANOVA). A subset of Hayden samples was randomly selected to match the number of samples in the Dartmouth cohort (n = 101). Vectors indicate the top three species responsible for separation of the two datasets based on Euclidean distance calculations. **C**, PCoA of Dartmouth and Hayden CF samples from ages 0-12 colored by presence of *Bacteroides* species (unweighted UniFrac, PERMANOVA). Samples containing no detectable *Bacteroides* are colored yellow and samples with > 0% relative abundance of *Bacteroides* are colored in blue. Vectors indicate the top three species responsible for separation of the two datasets based on Euclidean distance calculations. **D-E**, PCoA of Dartmouth and Hayden CF (purple) samples compared to TEDDY (**B**, orange) or DIABIMMUNE (**C**, green) samples from 0-12 months using unweighted UniFrac calculations. A subset of healthy samples in both cases were randomly selected to match the number of samples in the Dartmouth and Hayden CF combined group (n = 202). Vectors indicate the top three species responsible for separation of the two datasets based on Euclidean distance calculations. **F**, PCoA of DIABIMMUNE (green) samples compared to TEDDY (orange) samples from 0-36 months (unweighted Unifrac, PERMANOVA). A subset of TEDDY samples in was randomly selected to match the number of samples in the DIABIMMUNE group (n = 1128). Vectors indicate the top three species responsible for separation of the two datasets based on Euclidean distance calculations.

**Fig. S3.**
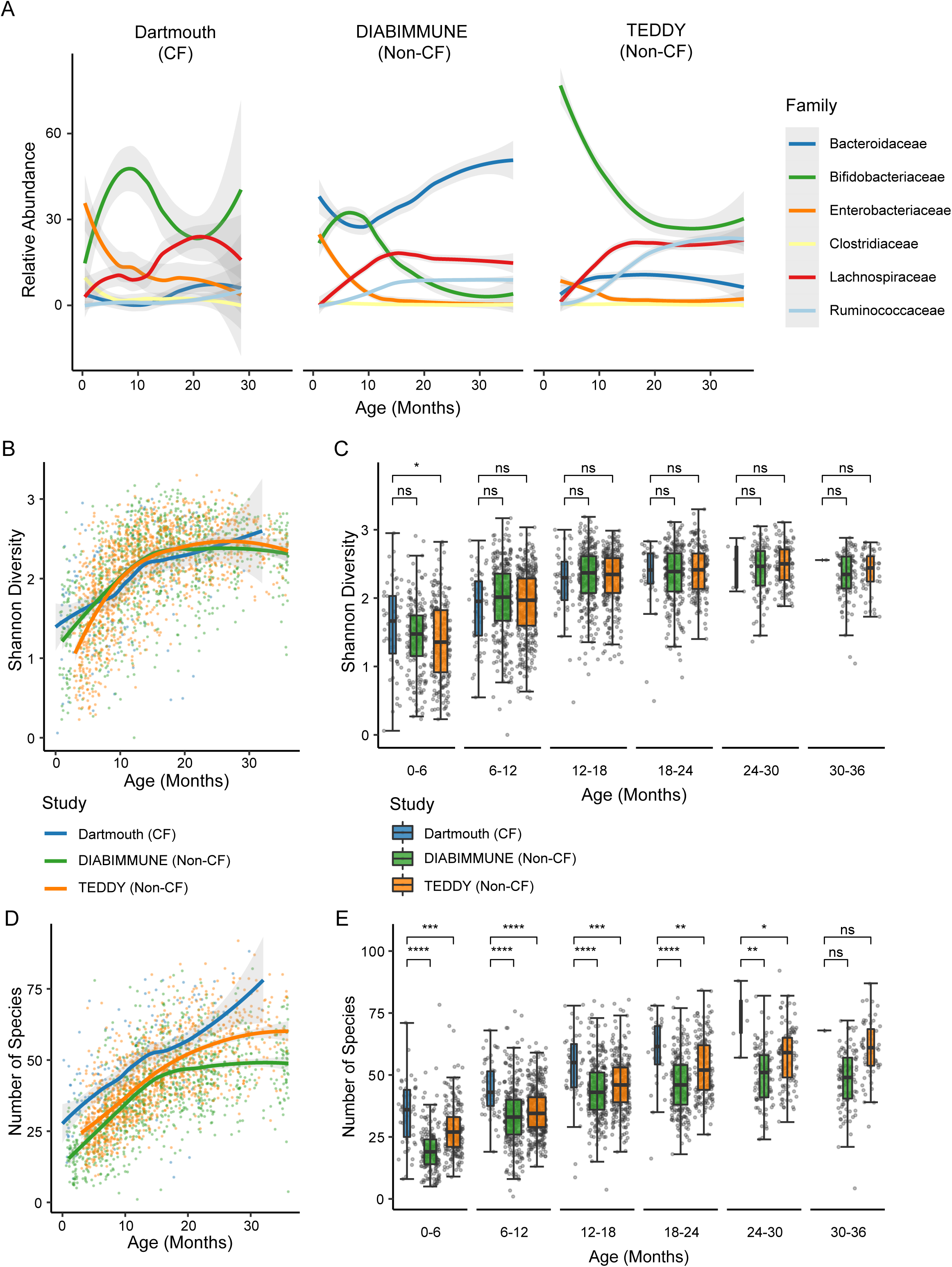
Altered microbiota compositional dynamics in CF infants. **A**, Relative abundance of select bacterial families in Dartmouth (left), DIABIMMUNE (center), and TEDDY (right) cohorts over time. Shaded regions signify 95% confidence intervals (n=2587). **B**, Shannon diversity plot of Dartmouth (blue), DIABIMMUNE (green), and TEDDY (orange) cohorts over time. Each sample is plotted as an individual datapoint and lines are plotted as averages over time. Shaded regions signify 95% confidence intervals (n=2589). **C**, Shannon diversity values of Dartmouth (blue), DIABIMMUNE (green), and TEDDY (orange) cohorts over time. Samples were binned into 6-month time bins, and statistical tests were run comparing Dartmouth to each of the non-CF control sets. The distribution of data is shown using a boxplot (n=2589). *P<0.05, Wilcoxon ranked sum test. **D**, Number of species in Dartmouth (blue), DIABIMMUNE (green), and TEDDY (orange) cohorts over time. Each sample is plotted as an individual datapoint, and lines are plotted as averages over time. Shaded regions signify 95% confidence intervals (n=2589). **E**, Boxplots of the number of species in Dartmouth (blue), DIABIMMUNE (green), and TEDDY (orange), over time. Samples were binned into 6-month time bins, and statistical tests were run comparing Dartmouth to each of the non-CF control sets (n=2589). *P<0.05, Wilcoxon ranked sum test.

**Fig. S4.**
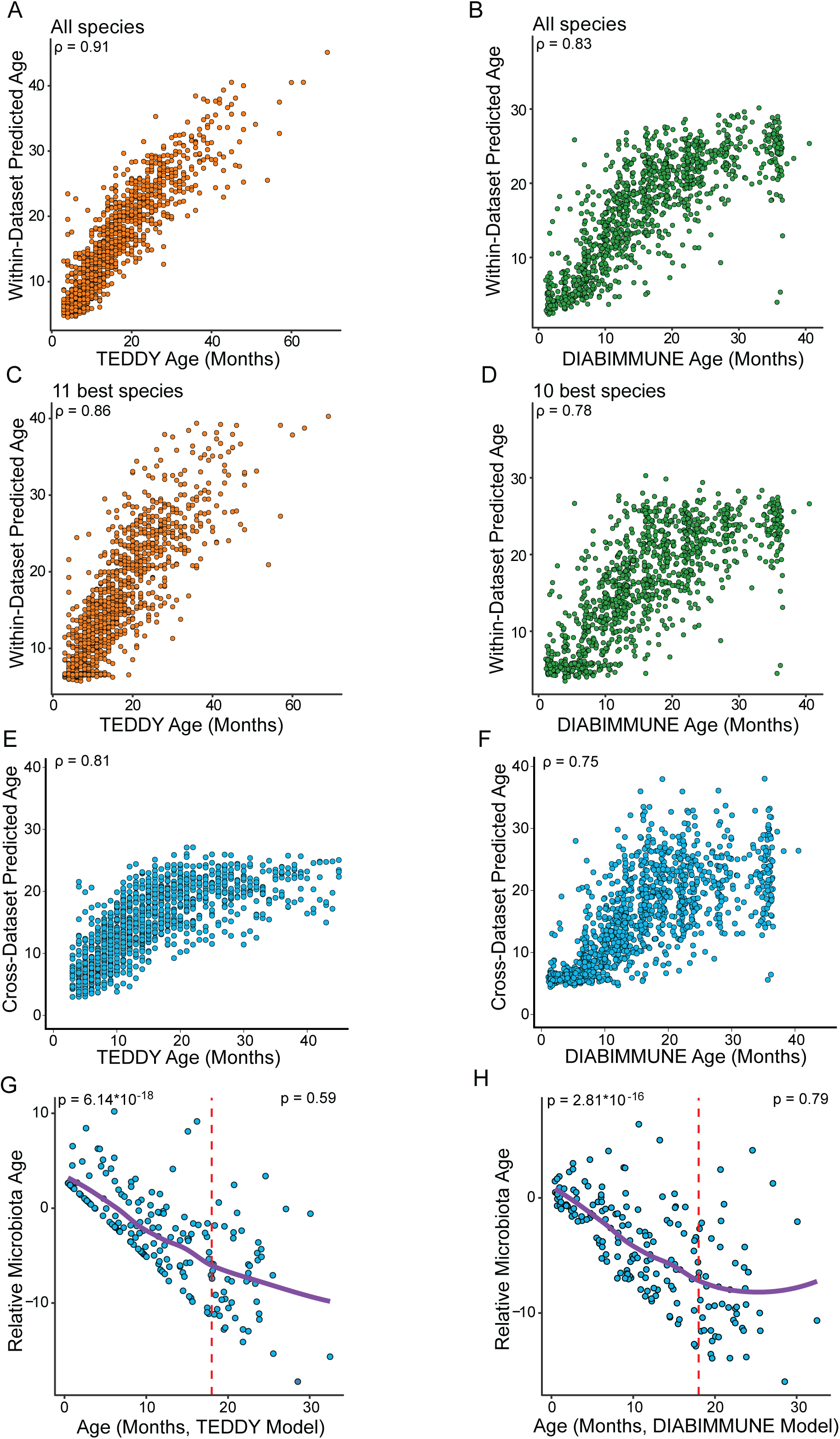
Random forest relative microbiota age model performance evaluated in training sets or cross-dataset. **A-B**, Scatter plots show the correlation between the random forest age model predicted age and the actual sample age. Two models were trained, one on non-CF samples from the TEDDY study (**A**, orange, n = 1246) and one on non-CF samples from the DIABIMMUNE study (**B**, green, n = 1154). **C-D**, Scatter plots show the correlation between the random forest age model predicted age and the actual sample age using the best 11 (**C,** orange, n = 1246, TEDDY cohort) or best 10 (**D**, green, n = 1154, DIABIMMUNE cohort) species. **E-F**, Within-dataset predictions were performed on the same training samples in 10-fold cross validation while cross-dataset predictions were done on the other dataset. Value in the upper left of each plot is the spearman correlation between predicted and actual age. **G-H**, Relative microbiota age of Dartmouth cohort when compared to the TEDDY (**G**, n = 190) or DIABIMMUNE (**H**, n = 190) age models using only the top 10 or 11 species. Red dotted line (18 months) signifies the time point at which the slope of the line is no longer significant. The p-values on the plots are a statistical test for a non-zero slope in a linear model between age and relative microbial age. P-value on upper left corner is slope prior to 18 months, p-value on upper right is slope after 18 months.

**Fig. S5.**
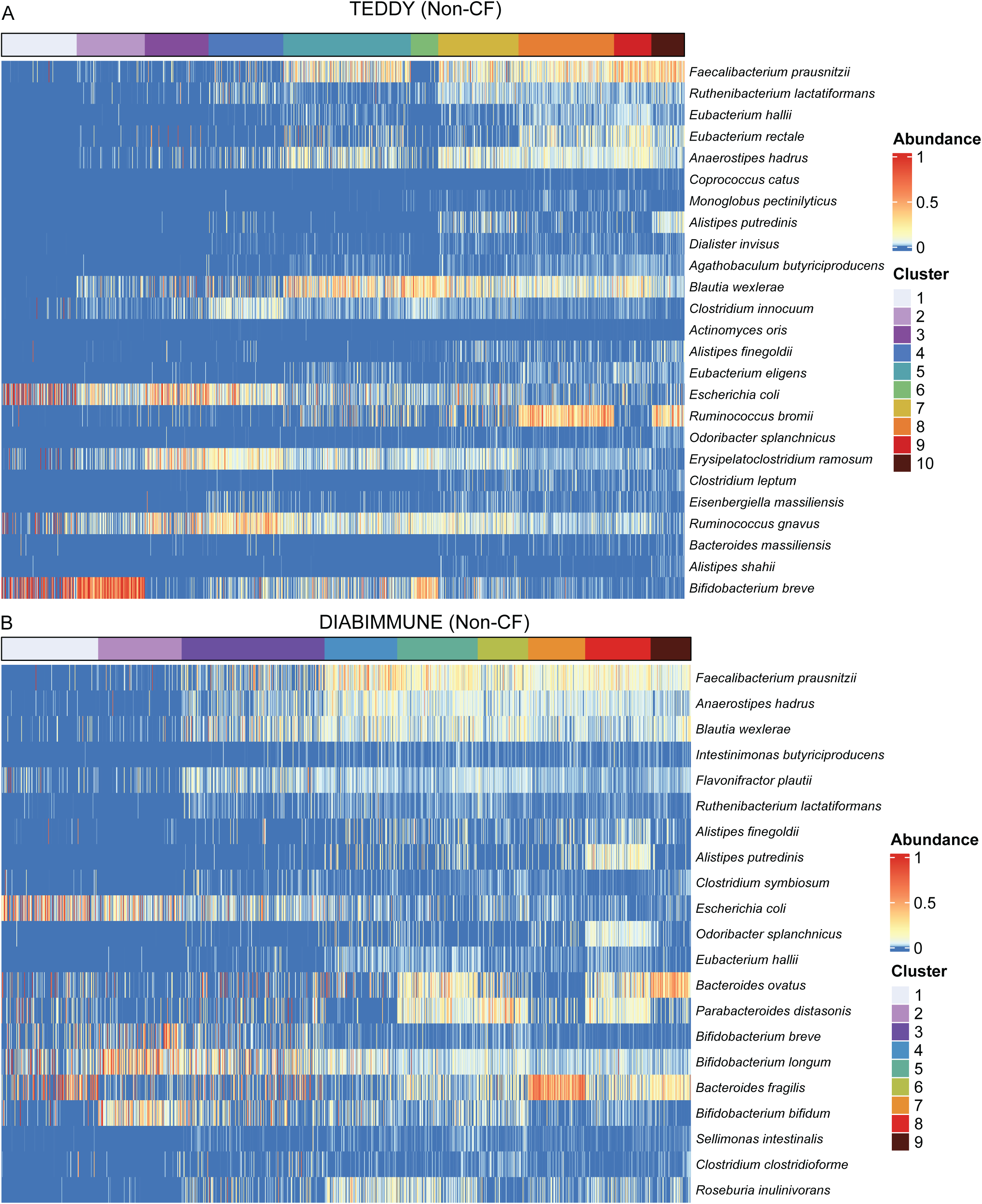
Heatmaps of species relative abundances in non-CF cohorts used in Dirichlet multinomial mixture modeling. **A**, Heatmaps of species relative abundance in different clusters derived from the Dirichlet multinomial mixture model. Shown here are all the top 25 species from the random forest age model that were used in the DMM clustering from the TEDDY non-CF cohort. Each column represents a different sample. Heatmap colors represent the relative abundance of each species. The bar at the top of the graph denotes which cluster samples were assigned to and are colored accordingly (n = 1246). **B**, Similar plot with data from the DIABIMMUNE cohort (21 species, n = 1154).

**Fig S6.**
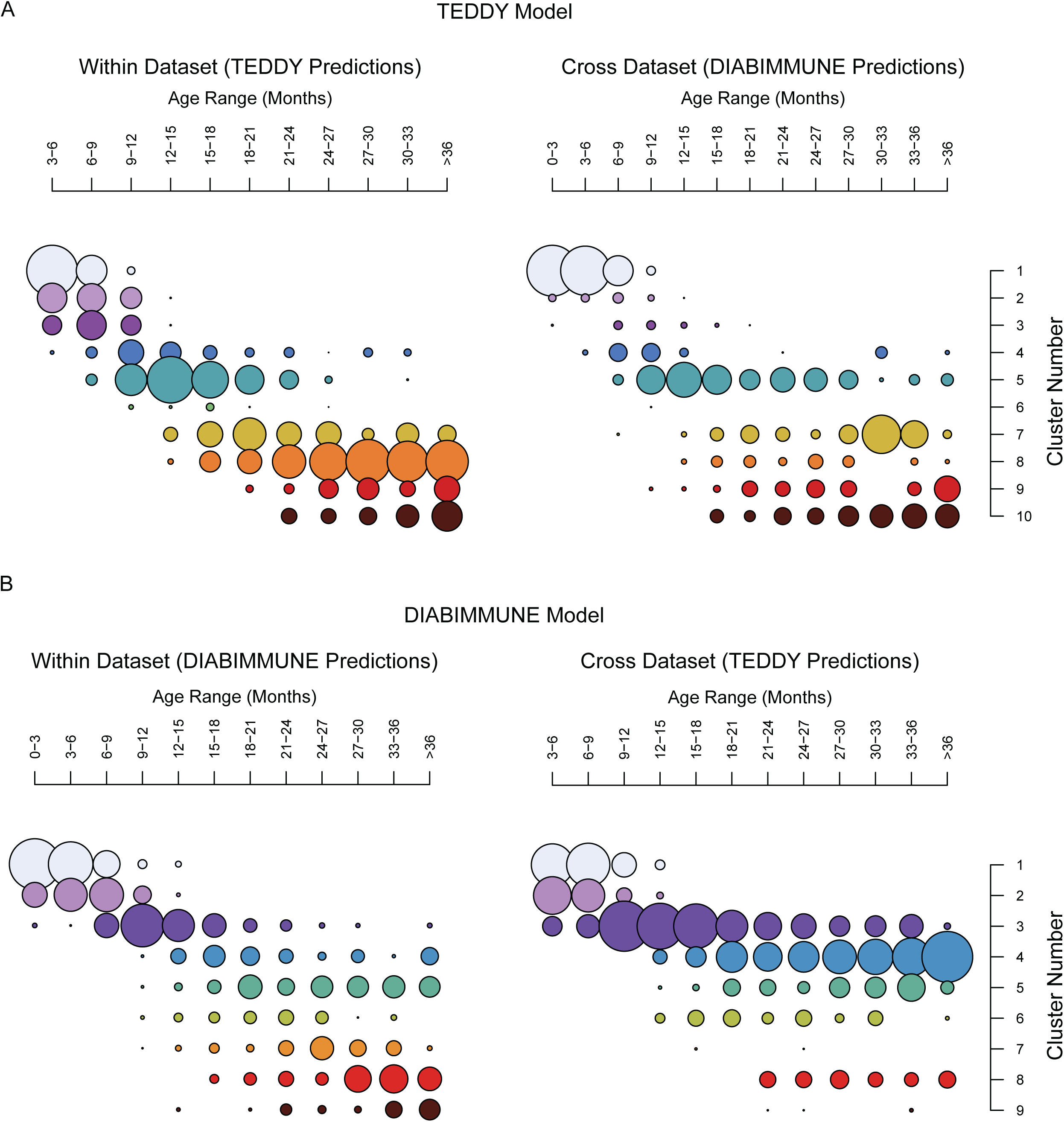
DMM cross-validation of non-CF cohorts shows consistent development of microbiome. **A**, Dirichlet multinomial mixture clustering (DMMs) of samples from the TEDDY (left), and DIABIMMUNE (right) cohorts using the TEDDY trained model (n = 2400). **B**, DMMs of infant samples from the DIABIMMUNE (left), and TEDDY (right) cohorts using the DIABIMMUNE trained model (n = 2400).

**Figure S7.**
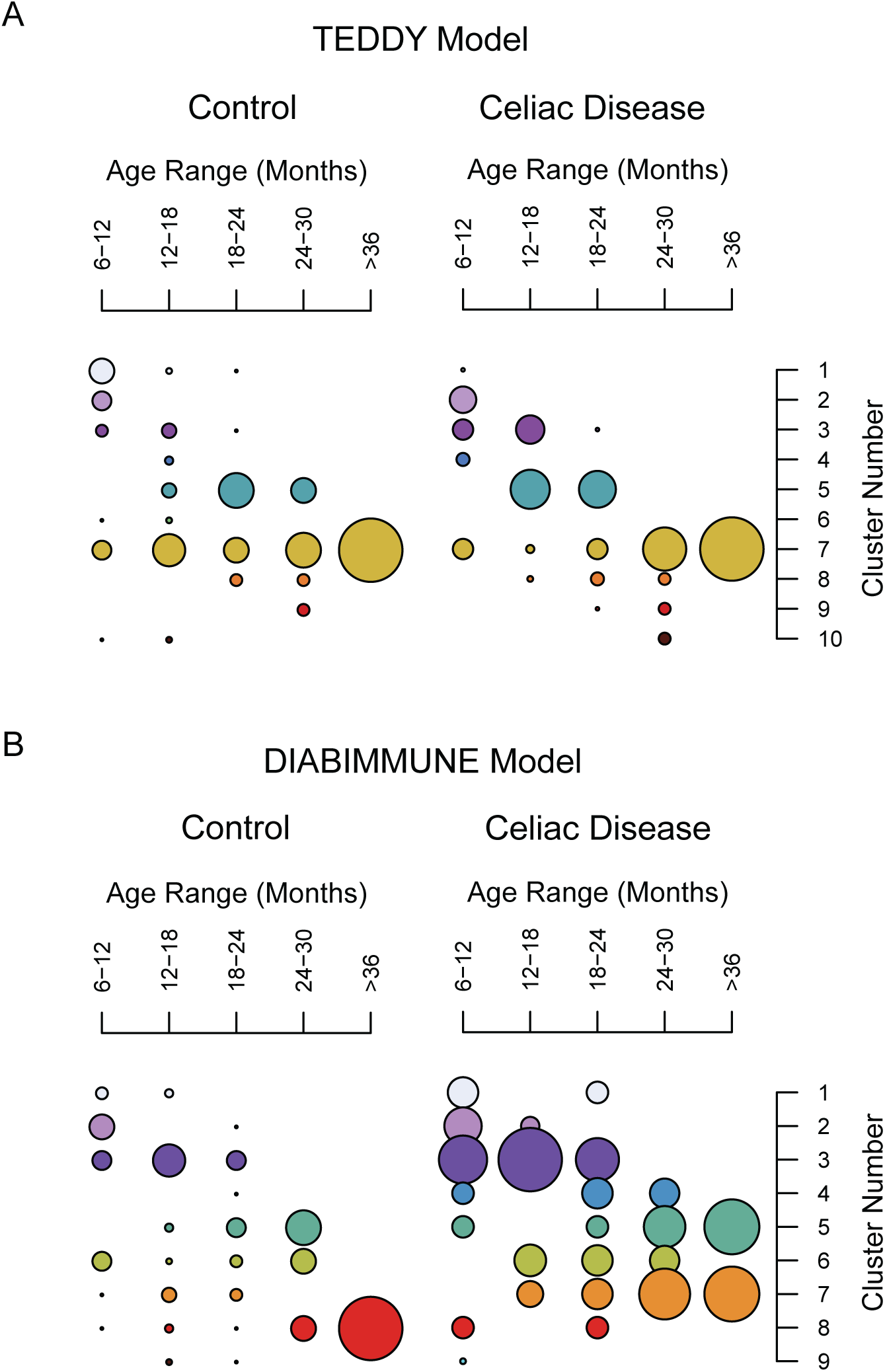
DMM modeling of gut microbiota from infants prior to onset of Celiac Disease. **A**, Dirichlet multinomial mixtures (DMMs) of infant samples from the Leonard study using the TEDDY trained model (n = 118). Control samples are found on the left and Celiac Disease (CD) samples are found on the right. Samples were binned into 6-month age bins and colors represent each cluster. **B**, DMMs of infant sample from the Leonard study using the DIABIMMUNE trained model. Leonard control samples are found on the left and Leonard CD samples are found on the right. Samples were binned into 6-month age bins and colors represent each cluster (n = 118).

**Figure S8.**
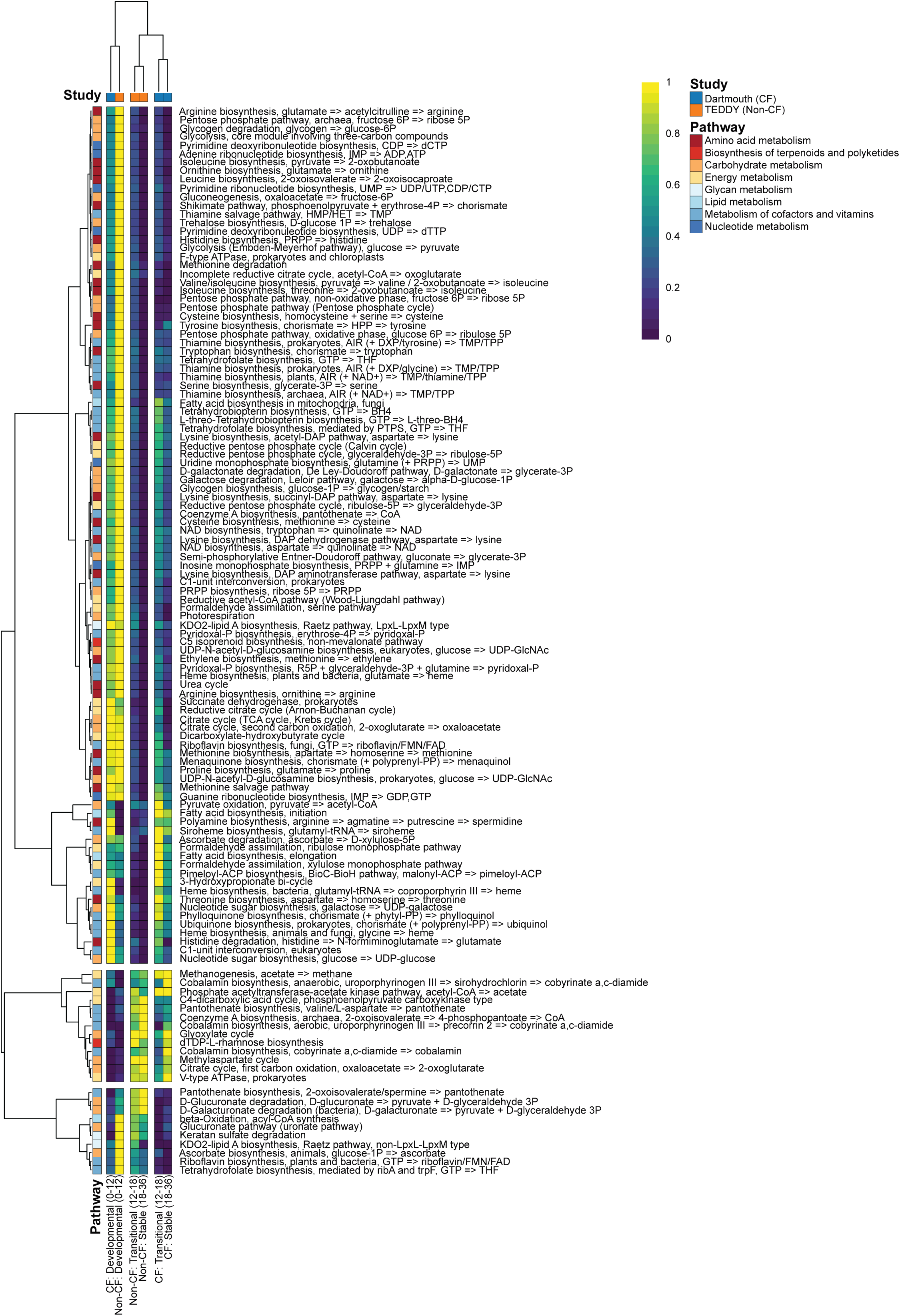
Functional gene abundance is altered in CF infant gut microbiota compared to TEDDY. Heatmap showing the mean abundance of all significant KEGG modules between the Dartmouth and TEDDY cohorts identified using MaAsLin at each developmental phase. Modules that were found to be significantly different in at least one phase are shown and values were normalized on a per row basis across all time points. The corresponding pathways for each module are also noted. (n=1436).

**Figure S9.**
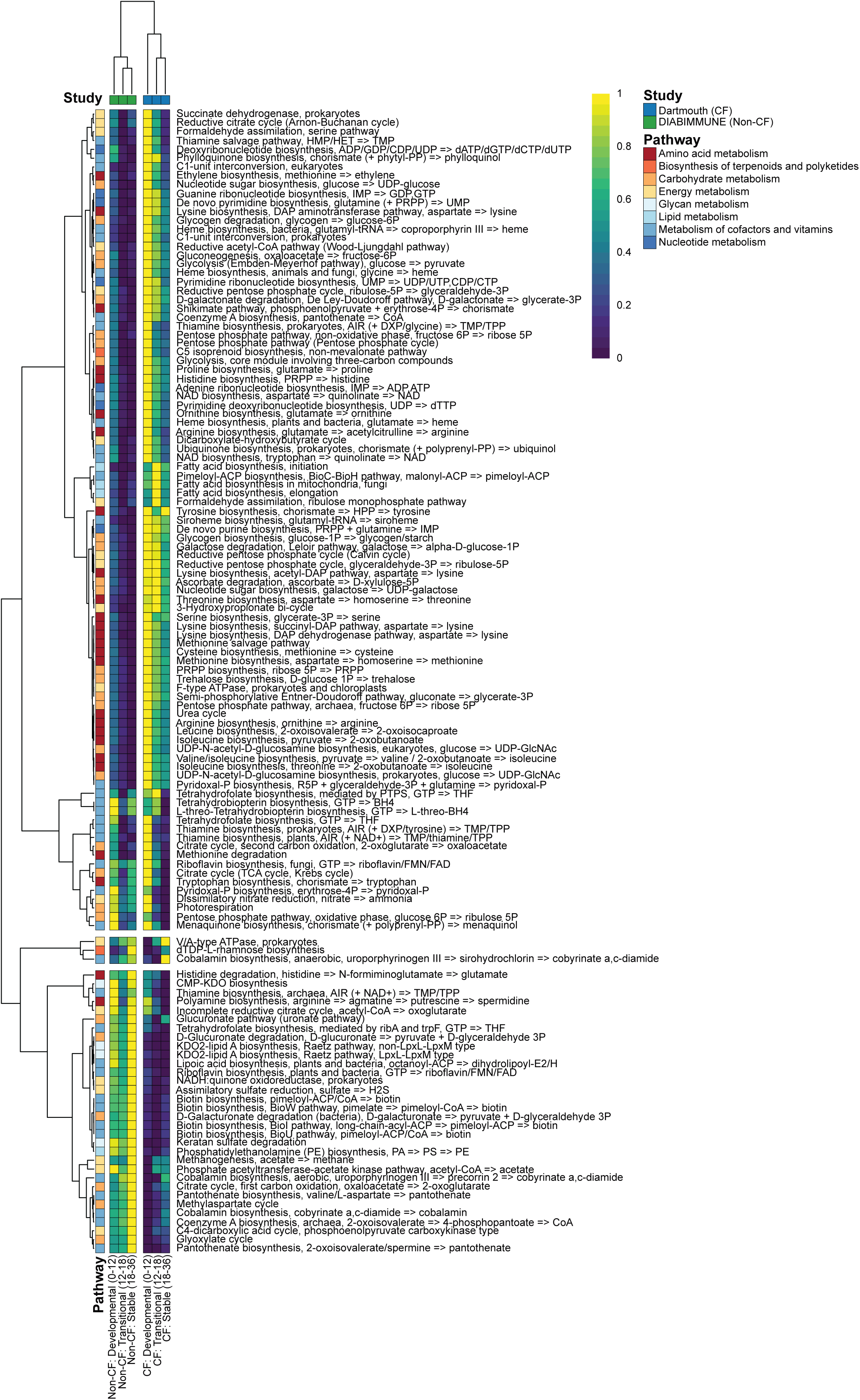
Functional gene abundance is altered in CF infant gut microbiota compared to DIABIMMUNE. Heatmap showing the mean abundance of all significant KEGG modules between the Dartmouth and DIABIMMUNE cohorts identified using MaAsLin at each developmental phase. Modules that were found to be significantly different in at least one phase are shown and values were normalized on a per row basis across all time points. The corresponding pathways for each module are also noted. (n=1344).

**Figure S10.**
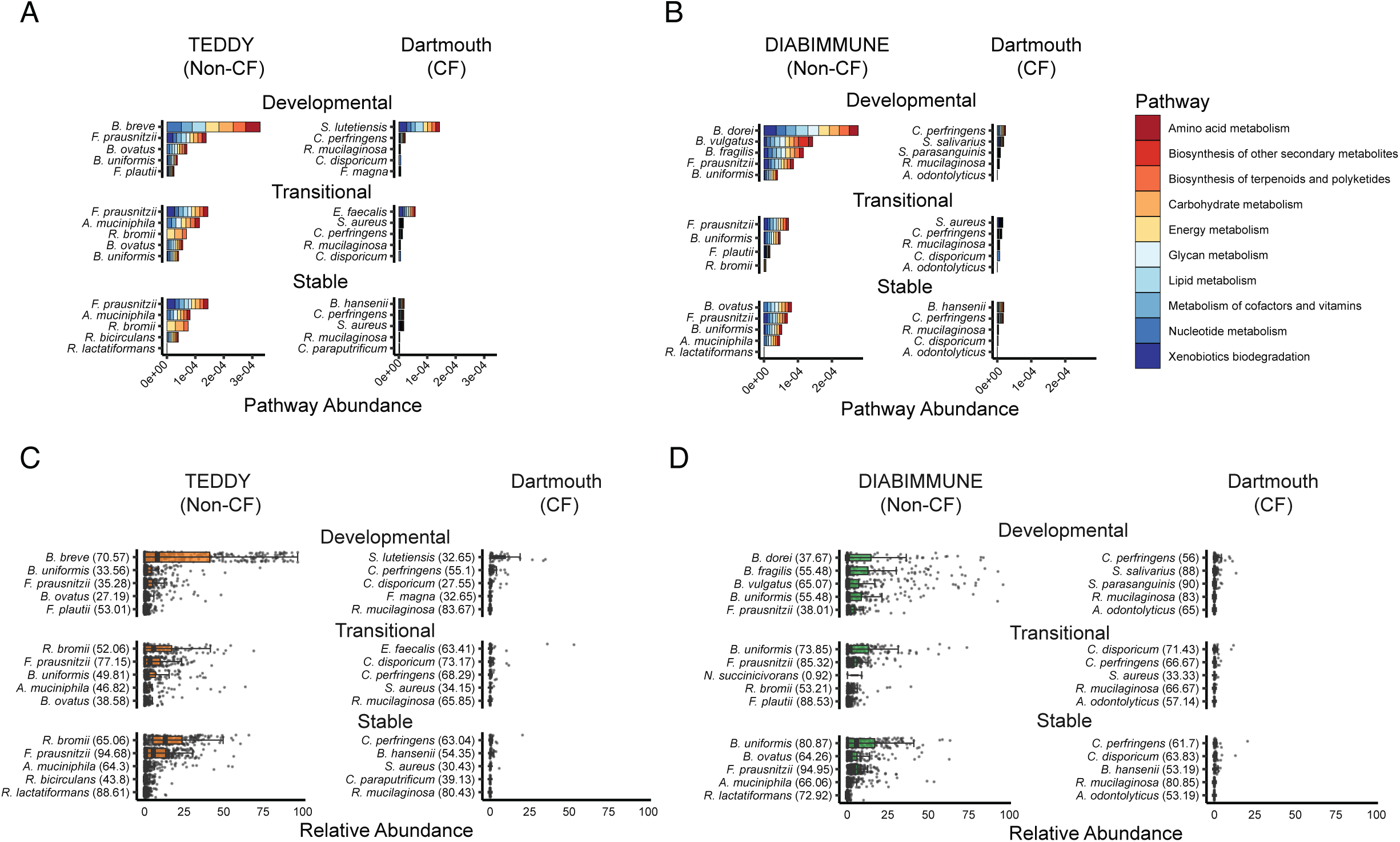
Altered functional capacity in CF compared to non-CF cohorts. **A-B**, Functional pathway abundances of the top five most statistically significant species at each developmental phase for the Dartmouth and TEDDY (**A**, n=1436) and Dartmouth and DIABIMMUNE (**B**, n=1344) cohorts. Significance was determined by Wilcoxon ranked sum test, and abundances were summed by KEGG pathway. **C-D**, Relative abundances of the top five most statistically significant species at each developmental phase for the Dartmouth and TEDDY (**C**, n=1436) and Dartmouth and DIABIMMUNE (**D**, n=1344) cohorts. Abundance was summed over all individuals for each phase, and the prevalence for each microbe is noted in parentheses.

**Figure S11.**
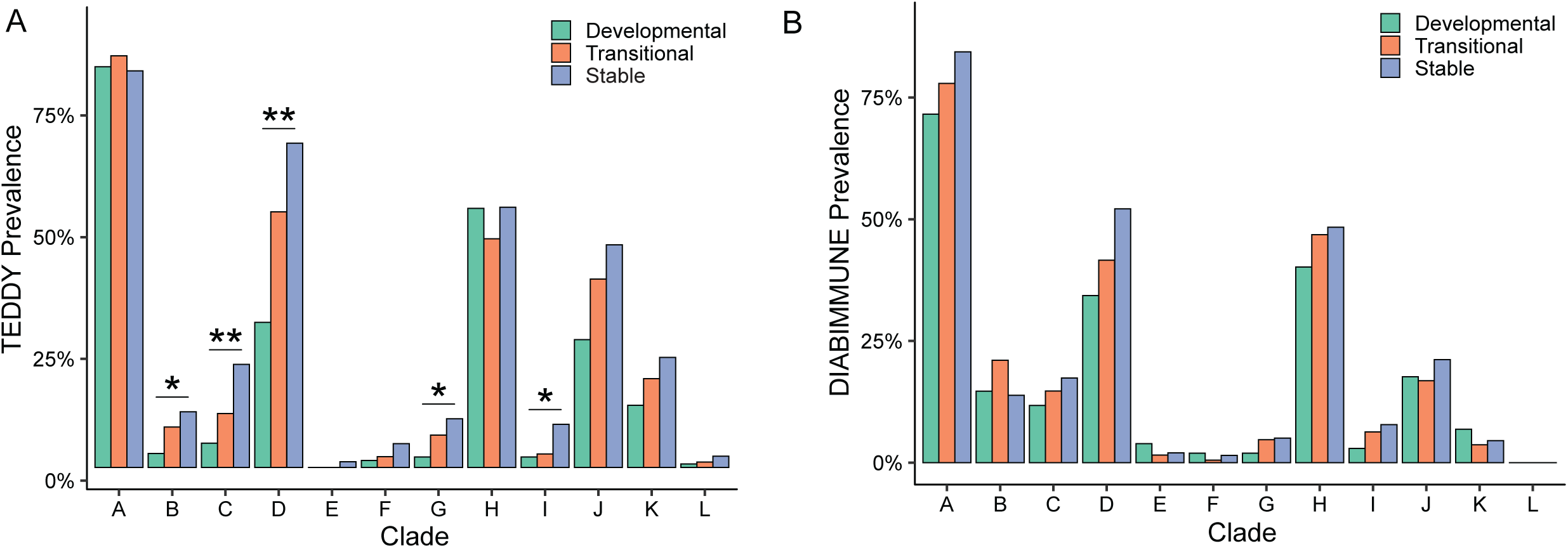
*F. prausnitzii* increases in prevalence over time in non-CF infants. **A**, Prevalence of each *F. prausnitzii* clade in the TEDDY cohort, colored by each of the developmental phases. *FDR <= 0.05, **FDR <= 0.01, Fisher’s exact test. P-values were adjusted for multiple corrections using an FDR (n = 1246). **B**, Prevalence of each *F. prausnitzii* clade in DIABIMMUNE cohort, colored by each of the developmental phases. *FDR <= 0.05, **FDR <= 0.01, Fisher’s exact test. P-values were adjusted for multiple corrections using an FDR (n = 1344).

## Supplemental Tables

**Supplemental Table 1 | Sample information for all datasets used in this study**

**Supplemental Table 2 | Species-level taxonomic profiling of all samples used in this study**

**Supplemental Table 3 | Age model species prevalence and relative abundances by developmental phase**

**Supplemental Table 4 | DMM cluster assignments for TEDDY control and Dartmouth CF**

**Supplemental Table 5 | DMM cluster assignments for DIABIMMUNE control and Dartmouth CF**

**Supplemental Table 6 | Additional sample information for select datasets used in this study**

**Supplemental Table 7 | Species-level taxonomic profiling of all samples used for Dirichlet Multinomial Mixture clustering**

**Supplemental Table 8 | Statistics for phase-level relative abundance comparisons between Dartmouth and TEDDY**

**Supplemental Table 9 | Statistics for phase-level relative abundance comparisons between Dartmouth and DIABIMMUNE**

**Supplemental Table 10 | Phase-level oral microbe fractions across datasets**

**Supplemental Table 11 | Total fungal prevalence across datasets**

**Supplemental Table 12 | Genus-level fungal prevalence across datasets**

**Supplemental Table 13 | Significant results from MaAsLin analysis between TEDDY and Dartmouth**

**Supplemental Table 14 | Significant results from MaAsLin analysis between DIABIMMUNE and Dartmouth**

## Notes

### Competing Interest Statement

The authors have declared no competing interest.

